# Flexible control of vocal timing in *Carollia perspicillata* bats enables escape from acoustic interference

**DOI:** 10.1101/2023.05.09.539989

**Authors:** Ava Kiai, Jan Clemens, Manfred Kössl, David Poeppel, Julio Hechavarría

## Abstract

In natural environments, background noise can degrade the integrity of acoustic signals, posing a problem for animals that rely on their vocalizations for communication and navigation. A simple behavioral strategy to combat acoustic interference would be to restrict call emissions to periods of low-amplitude or no noise. Using audio playback and computational tools for the automated detection of over 2.5 million vocalizations from groups of freely vocalizing bats, we show that bats (*Carollia perspicillata*) can dynamically adapt the timing of their calls to avoid acoustic jamming in both predictably and unpredictably patterned noise. This study demonstrates that bats spontaneously seek out temporal windows of opportunity for vocalizing in acoustically crowded environments, providing a mechanism for efficient echolocation and communication in cluttered acoustic landscapes.

**One Sentence Summary:** Bats avoid acoustic interference by rapidly adjusting the timing of vocalizations to the temporal pattern of varying noise.

## Introduction

The capacity for short-term vocal plasticity is advantageous in contexts where ambient noise is abundant, as it can enable acoustic jamming avoidance ^1^. Ambient noise presents a special challenge to echolocating bats, who rely on the returning echoes of their sonar pulses for navigation and in addition maintain social dynamics in part through the exchange of communication calls.

It is well established that bats possess impressive vocal plasticity, freely modifying various parameters of their vocalizations ^2^ such as the amplitude (known as the “Lombard effect”) ^3–7^, duration ^4,8–10^, repetition or emission pattern ^8,9,11^, complexity ^8^, and spectral content ^6,12,13^ (but see ^14–16^) in response to playback of interfering noise. Yet, how bats overcome interference from moment-to-moment fluctuations in the amplitude of continuous background noise, a situation analogous to their natural environment, has received less attention.

*Carollia perspicillata* bats live in colonies of up to hundreds of individuals where the acoustic landscape is densely populated by vocalizations which all share overlapping spectral and temporal properties. These bats emit highly stereotyped echolocation pulses comprised of brief (∽1-2 ms) ^17^, multi-harmonic, frequency-modulated downward sweeps (peak frequency 60-90 kHz) ^18,19^. This species also possesses a repertoire of social calls, some of which have been associated with specific behaviors, such as distress, territorial aggression (males), courtship (males), and the eliciting of maternal attention (infants) ^20–22^. These communication calls typically feature multiple distinct harmonics, with the most energy concentrated in lower frequencies (below 50 kHz) ^20^.

In this study, we investigated the ability of bats to adapt the timing of their vocalizations (both echolocation and communication calls) to overcome acoustic jamming, using temporally predictable and unpredictable noise, across two experiments. We hypothesized that bats would preferentially vocalize in periods of low amplitude in amplitude modulated noise, in line with a metabolically efficient signal optimization strategy. Humans regularly employ a similar strategy, such as when a pair of speakers pause their conversation so as not to be drowned out by the blaring siren of a passing ambulance.

We observed that freely vocalizing bats flexibly adapt the timing and rate of their calling to be inversely proportional to dynamically-changing background amplitude levels. This temporal jamming avoidance behavior emerged in the presence of both predictably and unpredictably patterned noise, implying an underlying auditory-vocal circuit that does not require entrainment for optimizing call timing. In addition, calling behavior is modulated not only by instantaneous amplitude levels but also by more global sound statistics (i.e., second-order temporal patterns), suggesting that bats learn and exploit properties of the acoustic environment which unfold over time.

## Results

### Bats cluster call onsets toward amplitude troughs in broadband masking noise

In *experiment 1*, we recorded vocalizations from eight groups of six bats during a silent baseline and during playback of two types of white noise featuring different carrier frequencies (a 10-96 kHz “broadband masker”, which overlaps with both communication and echolocation call frequencies, and a 50-96 kHz “high-frequency masker” (hereafter “high-freq masker”), which overlaps in frequency only with echolocation calls) (Fig. 1A-B). Audio recordings from our colony of captive bats showed that spontaneous vocalizations feature a prominent rhythm at approximately 11 Hz (Fig. S1). Thus, we amplitude modulated the two maskers at 8 Hz and 15 Hz to see if the bats could adjust to slower or faster rates, respectively (Fig. 1C).

**Fig. 1.**
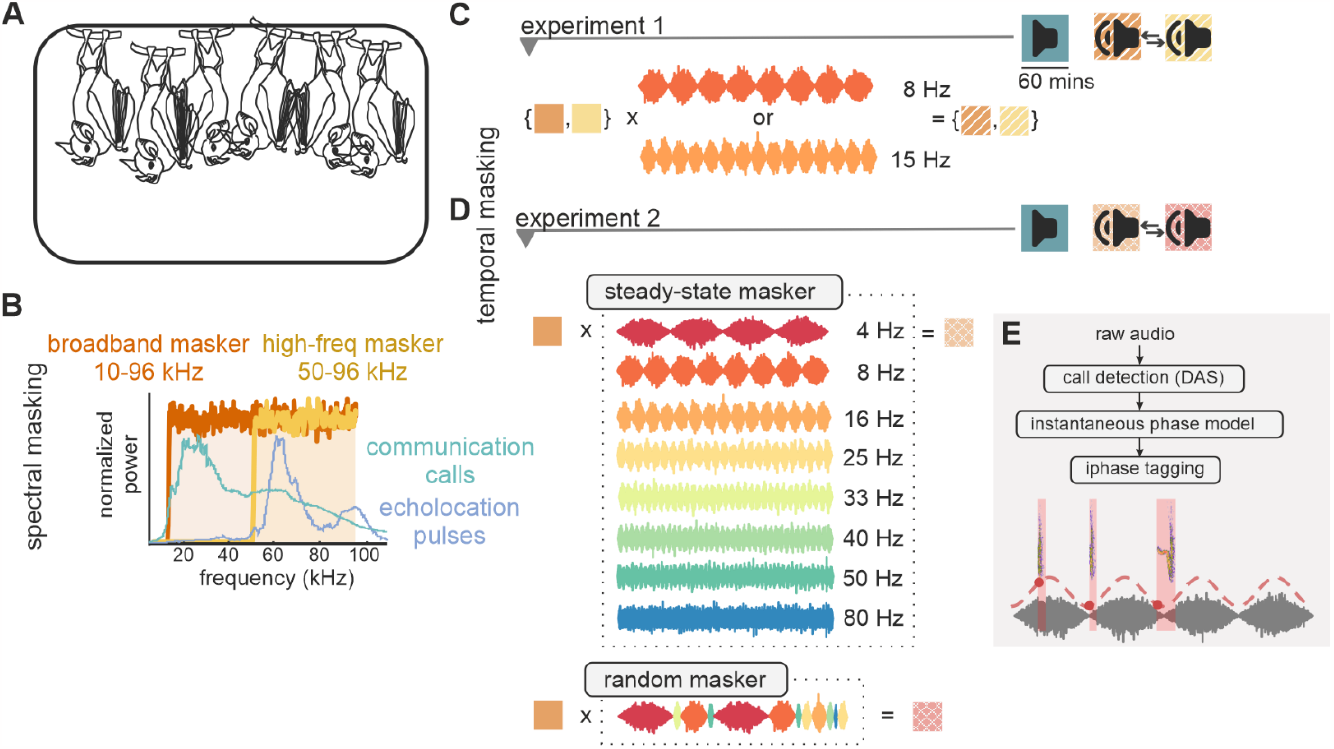
Schematic of experiments. **(A)** Each group of bats consisted of 6 adults (4 male) that could flit and socialize freely in a cage placed within the recording chamber. (**B)** Stimuli were broadband white noise with 10-96 kHz (broadband masker) and 50-96 kHz (high-freq masker) carrier frequencies. Teal and violet traces indicate normalized power spectra of *C. perspicillata* communication and echolocation calls, respectively. (**C)** In *experiment 1*, maskers were amplitude modulated at 8 or 15 Hz for each group. Procedure (right): Recording days (5) consisting of three one-hour blocks: a silent baseline, then playback of the broadband and high-freq masking noise, counterbalanced. (**D)** In *experiment 2*, broadband masking noise was amplitude modulated at eight amplitude modulation (AM) rates (4-80Hz). A random condition consisted of a randomly permuted sequence of the eight AM cycles. Procedure (right): Recording days (5) consisted of a silent baseline, then playback of the steady-state maskers (playback of each modulation rate for 7.5 minutes each in randomized order), and random masking noise, counterbalanced. (**E)** Data analysis: Call events (pink shaded areas) were detected using Deep Audio Segmenter (DAS). Calls were tagged with the instantaneous phase (red dots) of the amplitude envelope (red dashed line) at call onset time.

We labelled detected vocalization onsets with the instantaneous phase (0 to 2π) of the modulation cycle at the corresponding time point (Fig. 1E). For the silent baseline, we labelled vocalizations according to a cosine model of a fictitious amplitude modulation with the same rate as the corresponding masking noise. Based on visual inspection of a subsample of our data (Table S1), and the fact that these calls were primarily short in duration (median = 3.4 *ms*, IQR= 3.3 *ms*, 75% of calls < 5 *ms*, across both experiments), we estimate that most (∽ 90%) of detected vocalizations were echolocation pulses.

As the nature of these call onset data are cyclical (i.e. calls occurring at the end of an amplitude modulation cycle may also be considered as occurring at the start of the following cycle) (Fig. 2A), we represented the distribution of call onsets in the polar as well the cartesian plane (Fig. 2B). We also made use of statistics for the analysis of circular data (see Methods) which take into consideration the temporal proximity of values that fall at the boundary of consecutive modulation cycles.

**Fig. 2.**
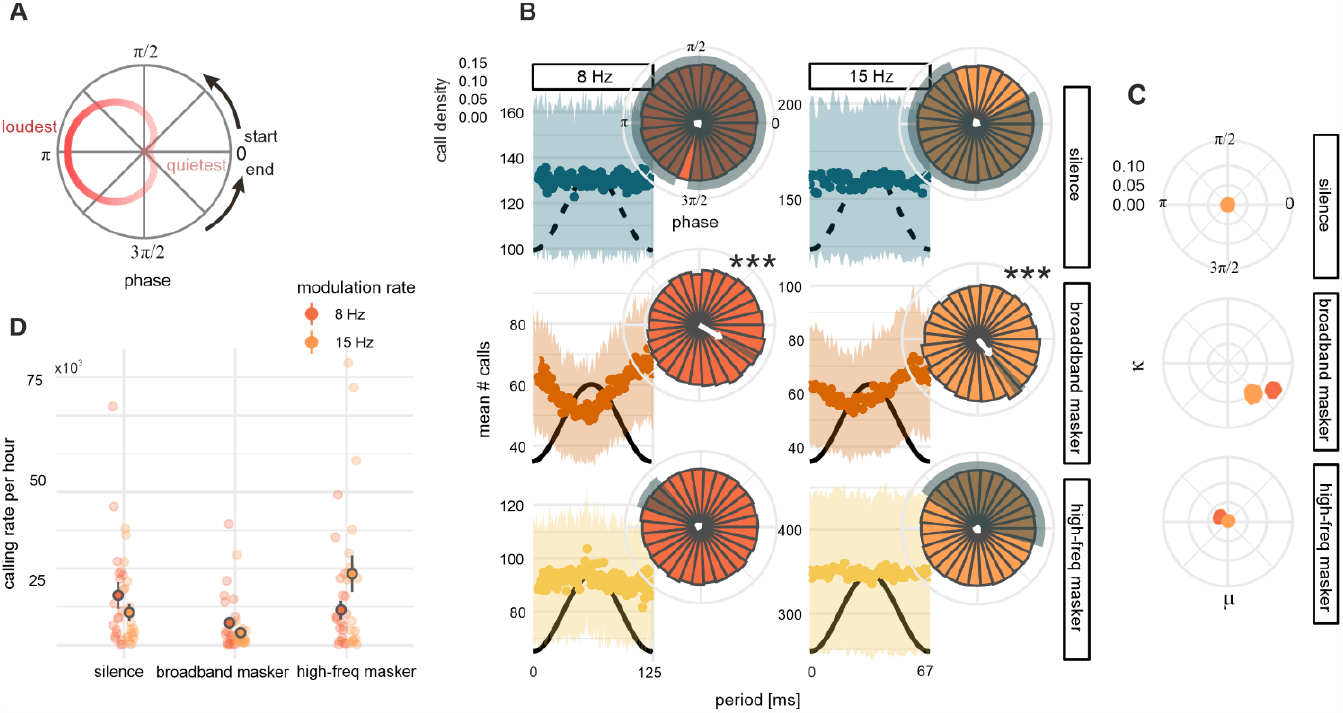
Anti-phase preference for call onset timing in broadband noise. (**A)** Schematic representing the translation from the cartesian to polar coordinates. 0 phase (right) corresponds to the amplitude trough. Red trace represents amplitude change over a cycle. (**B)** Average call onset distributions within the modulation cycle track the inverse of the amplitude phase in broadband masker conditions, but not in silent and high-freq masker conditions (1 ms bins). Shaded areas indicate standard error of the mean (SEM). Black curves: **schematic** of amplitude envelope (not to scale) in playback conditions and simulated amplitude envelope in silent conditions. Insets: Circular density histograms of call onsets (30 bins). White arrows indicate resultant vectors: arrows indicate angular means; arrow lengths indicate resultant lengths (concentration of data at the angular mean). Shaded areas indicate maximum likelihood (MLE) 95% confidence intervals for angular means. Stars indicate significant p-values from Rayleigh ‘s test of uniformity. *** *p < 0*.*001* (**C)** Bootstrapped MLE von Mises mean (μ) and concentration (κ) parameters indicating robust pattern of call onset clustering in the broadband masking condition for both modulation rates. (**D)** Data points (lightly colored dots) indicate number of calls observed per hour for each group for each recording day. Estimated mean calling rates (darkly colored dots with gray outline) from negative binomial models show a drop in calling between silent baseline and acoustic masking conditions (except for 15 Hz high-freq masker condition). Gray vertical lines indicate SE of model fit for predicted means.

We found that bats preferentially vocalized in the quieter phases of the ongoing amplitude modulation noise, rendering the distribution of call onsets within the cycle inversely proportional to the amplitude level of the playback noise (Fig. 2B). Call onset density distributions were strongly unimodally clustered toward the amplitude downstate (Fig. 2B, middle row) in the presence of the broadband masking noise for both modulation rates (Rayleigh ‘s test: 8 Hz: = *r* 0.06, *p* < 0.001; 15 Hz: = *r* 0.03,*p* < 0.001, Bonferroni adjusted). However, call onsets emitted in the high-freq masking noise were not strongly clustered at any particular phase in the modulation cycle, more closely approximating a uniform circular distribution (8 Hz: *r* = −0.001,*p* = 1; 15 Hz: = *r* 0.001,*p* = 1, Fig. 2B, bottom row). Importantly, no bias towards vocalizing at either rate was observed in the silent baseline (8 Hz: *r* = −0.001,*p* = 1; 15 Hz: *r* < = −0.001, = 1) for either modulation rate (Fig. 2B, top row). This preference for calling in the downstate of the amplitude cycle in the broadband masking condition was present for all groups tested (4 for each modulation rate) and on all five recording days (Fig. S2).

To concisely describe the distribution of call onsets in the modulation cycle, we treated data points (phase values at which call onsets occurred) as unit vectors and computed their sum (the resultant vector). Resultant vectors indicate both the direction (phase) at which the mean is located (Fig. 2B, white arrow tips) and the degree to which the data are concentrated at that direction (Fig. 2B, arrow lengths). A resultant vector length of 0 would indicate that the data are uniformly spread along the circle, while a length of 1 would indicate that all data points occupy the same location. Resultant vectors for calls emitted in broadband masking conditions indicated that call onsets were prominently clustered near amplitude troughs (Fig. 2B, Table S2).

To confirm that this result is robust feature of the data, we computed maximum likelihood von Mises parameters, the circular mean (μ) and concentration (κ), in a bootstrap procedure. These parameters revealed that the clustering of call onsets on the falling edge of the amplitude cycle (Fig. 2C, lower right quadrants) was consistent throughout the dataset for both modulation rates, but only in the broadband noise condition (Fig. S3A).

Phases at which call onsets occurred varied significantly between playback conditions for each modulation rate (Mardia-Watson-Wheeler test: 8 Hz: 1228, *p* < 0.001; 15 Hz: 369, *p* < 0.001, Bonferroni adjusted). The spread of call onsets (angular dispersions), but not the angular means, were significantly modulated by the type of masking noise (Rao ‘s test: broadband vs. high-freq maskers, *p* < 0.001, Table S3).

### Rate of calling is modulated by the degree of spectral masking

Playback noise impacted not only the timing, but also the number of vocalizations emitted by the bats. The presence of masking noise resulted in a reduction in the rate of vocalization between silent (528,155 calls) and broadband masking conditions (224,384). Surprisingly, the rate of calling increased relative to the silent baseline in the presence of the high-freq masker (672,528).

Playback condition significantly accounted for the variation in the hourly rate of calling for the 15 Hz context (*χ*^2^ = 21.34(2),*p* < 0.001), but not in the 8 Hz context (*χ*^2^ = 4.37(2), *p* = 0.11), as modelled by a negative binomial distribution. Nonetheless, in the 8 Hz context, the rate of calling dropped between silent and broadband masking conditions (B = .45, SEB = 0.38, *p* = 0.034 95% CI[0.21 − 0.95]), and silent and high-freq masking conditions. In the 15 Hz context, the rate of calling dropped between silent and broadband masking conditions (B = .39, SEB = 0.36,*p* < 0.01, 95% CI[0.19 − 0.79]), but increased between baseline and the high-freq masker (B = 2.19, SEB = 0.36,*p* = 0.03, 95% CI[1.07 − 4.46], Table S4-7, Fig. 2D). Between modulation rate contexts, calling rates were only significantly different in the high-freq masking condition (*z* = −1.85,*p* = 0.06), due to the greater number of calls in the 15 Hz context.

### Bats can adapt call timings to both predictably and unpredictably patterned noise

In *experiment 1*, we observed that bats exhibit an untrained and flexible adaptation of vocalization timing and rate when presented with rhythmic masking noise. In *experiment 2*, we further probed this behavior by asking: First, what is the upper temporal limit for this anti-phase calling behavior? And second, can the bats still perform this feat if the temporal pattern of the masking noise is unpredictable? To this end, we played the broadband masker noise to four additional groups of six bats, this time featuring amplitude modulation at eight different rates (4, 8, 16, 25, 33, 40, 50, and 80 Hz) for 7.5 minutes each (steady-state condition). To answer the latter question, we also generated a masking noise with a randomly permuted sequence of amplitude modulation cycles sampled from those eight rates for 60 minutes (random condition) (Fig. 1D).

Call onsets tracked the inverse of the modulation envelope up to 16 Hz (Rayleigh ‘s test: 4, 8 and 16 Hz, *p* < 0.001, Bonferroni adjusted, Fig. 3A). Call onset clustering was negligible for rates of 25 Hz and above (Table S8). Importantly, as this anti-phase clustering pattern was present in both steady-state and random temporal conditions, the bats evidently did not need to be able to predict the time-of-arrival of the upcoming amplitude downstate to be able to adapt call timings (Fig. 3A). This call pattern was present for all groups of bats tested (Fig. S4).

**Fig. 3.**
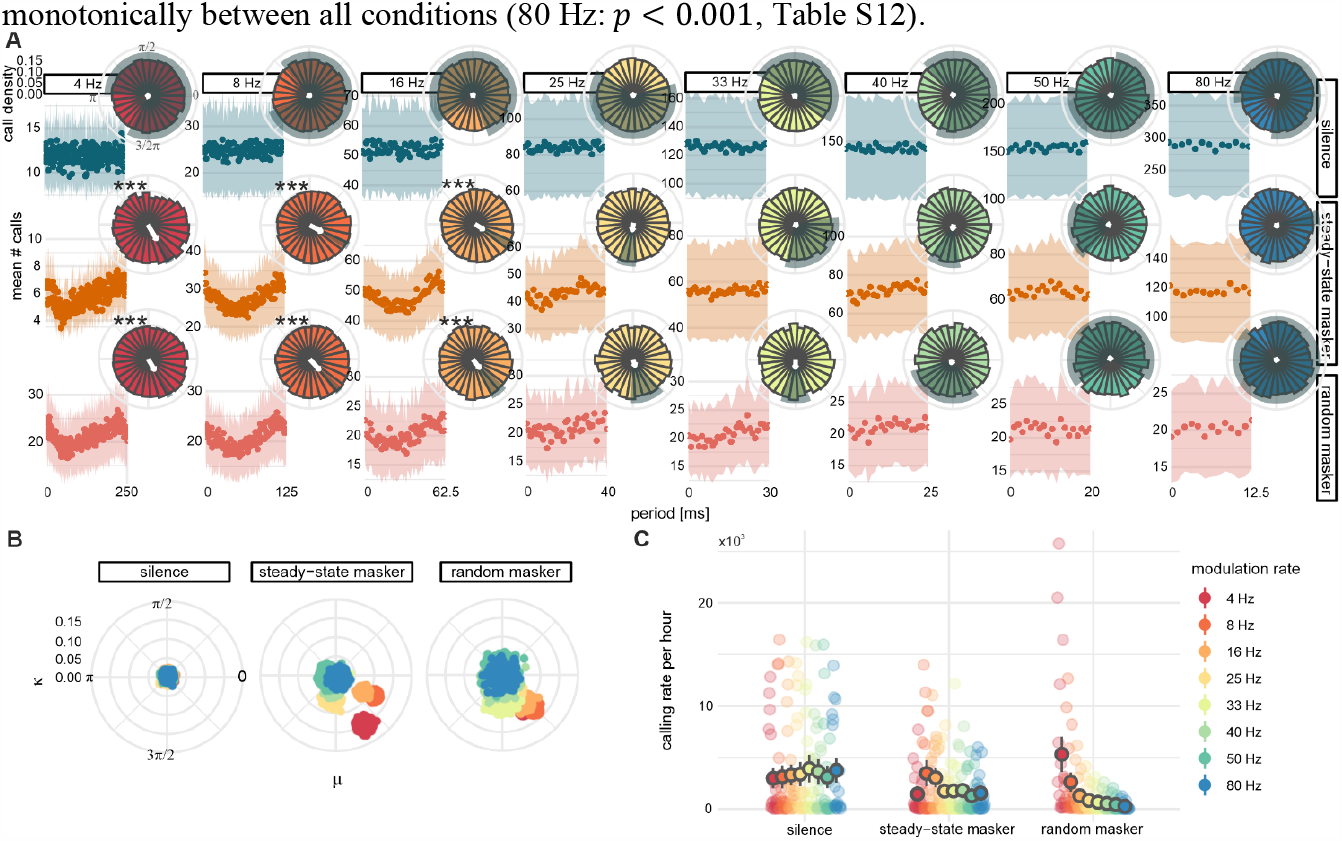
Call timing adaptation to amplitude modulated noise is independent of predictability but sensitive to rate. (**A)** Average call onset distributions show call timings follow the inverse of the amplitude modulation cycle in both predictable (steady-state) and unpredictable (random) temporal contexts, but only do so reliably up to 16 Hz. *** *p < 0*.*001*, Rayleigh ‘s test. (**B)** MLE von Mises mean and concentration parameters indicate that the call clustering pattern is robust but graded in playback conditions, with the greatest anti-phase concentration of call onsets occurring in slowest rate contexts and the clustering becoming less extreme with increasing rate. (**C)** Estimated mean rates of calling (darkly colored dots with gray outline) from negative binomial models show suppression of calling rate induced by masking noise, but the degree of suppression is determined by modulation rate. Data points (lightly colored dots) indicate number of calls observed per hour for each group for each recording day. Gray vertical lines indicate SE of model fit for predicted means.

Notably, bootstrapped von Mises parameters showed that this temporal “targeting” of the falling edge of the amplitude modulation does not display a step change above 16 Hz, but rather a gradual decrease in tracking fidelity (Fig. 3B, Fig. S3B).

Phases at which call onsets occurred varied significantly between playback conditions for modulation rates from 4 to 40 Hz (Mardia-Watson-Wheeler test: *p* < 0.001, Bonferroni adjusted, Table S9). Angular dispersions were significantly modulated by playback condition for 4 to 25 Hz (Rao ‘s test: *p* < 0.001) and more modestly in the 33 and 40 Hz contexts (*p* < 0.05). Angular means were only markedly different between playback conditions for the 8 Hz context (*p* < 0.001, Table S10).

### Rate of calling depends on local and global acoustic context

In line with our hypothesis that “noisier” acoustic environments incur greater suppression of vocalization, most calls detected in *experiment 2* were emitted in the silent condition (539,086), with fewer calls emitted in the presence of the steady-state masker (312,728), and the fewest calls emitted during playback of the random masker (237,180). However, the precise pattern of suppression was sensitive to the temporal structure of the acoustic masker.

Most notably, while playback of the random masker reduced the overall number of vocalizations, there was significant variation in the rate of calling observed in cycles of each modulation rate in this condition (*χ*^2^ = 81.08(7), *p* < 0.001, Fig. 3B, Table S11). More calls were observed in 4 Hz cycles when those cycles were embedded in the random stream of amplitude modulations than when playback consisted of only a continuous stream of 4 Hz cycles (*p* = 0.01, Fig. 3B, Table S12). This finding may be because the unpredictable stream posed a significant challenge to the bats which could be partially overcome by exploiting the comparably slow sound level rise and decay, extended over 250 ms, provided by the 4 Hz cycles.

Meanwhile, vocalization rates in 8 and 16 Hz contexts were comparable across all conditions (Fig. 3B, Table S12-14), possibly due to the relative ease of shifting call timing at rates close to the spontaneous ∽ 11 Hz vocalization rate.

For all other modulation rates, playback condition was a significant predictor of the variance in the number of observed vocalizations (*p* < 0.05, Fig. 3B, Table S13-14), which dropped significantly between the silent baseline and random masker conditions (25 - 50 Hz: *p* ≤ 0.01, Table S12) or monotonically between all conditions (80 Hz: *p* < 0.001, Table S12).

### Narrowing temporal windows of opportunity for vocalization leads paradoxically to fewer overlapping calls

In modulated noise, temporal windows of opportunity for vocalizing are sparse. If groups of bats begin collectively targeting narrow windows for vocalizing, this could lead to an increase in the number of temporally overlapping calls. Although our study design did not permit an evaluation of individual calling patterns, temporal overlaps in detected calls nonetheless signified multiple speakers. Overall, we found few overlapping calls (*experiment 1*: 42,618 of 1,425,067 calls, *experiment 2*: 31,942 of 1,088,994; < 3% in total; Fig. S5B, D, Table S15).

However, contrary to our predictions, the fewest number of overlaps were recorded in the masking conditions where acoustic interference was greatest and would have encouraged the greatest temporal clustering of calls (Fig. S5B, D, Table S16). Nonetheless, overlapping calls were clustered in the amplitude downstate for slower modulation rates (Rayleigh ‘s test: experiment 1: 8 Hz broadband masker: *p* < 0.001, 15 Hz high-freq masker: *p* < 0.001; experiment 2: 8 Hz steady-state masker, *p* = 0.003, 16 Hz steady-state masker, *p* = 0.04, Bonferroni adjusted; Fig. S5A, C).

### Evidence for temporal anchoring to terminal troughs across different temporal rates

The fact that vocal timing can be calibrated to occur in an anti-phase pattern within a single amplitude modulation cycle (Fig. 3A, random masker) implies that acoustic evidence in the first half of the cycle (the rising edge) is sufficient to inform the bats’ decision of when to vocalize in the second half of the cycle (the falling edge). Yet, the rates for which we observed this adaptation (4, 8, 15 and 16 Hz) feature significantly different period lengths (250 to 62.5 ms), leaving open the question of whether bats achieve this timing adaptation by attempting to call after amplitude peaks or by targeting the terminal troughs.

To answer this question, we computed two measures of call timing from bootstrapped mean call onsets (in radians): time relative to the amplitude peak, and time relative to the terminal amplitude trough (in ms) (Fig. 4A, C). If amplitude peaks are used as acoustic landmarks for timing adaptation, then calls should arrive at roughly the same delay after the peak, independently of rate. Alternatively, if terminal troughs are targeted, then calls should arrive at similar delays before the trough, across different rates.

**Fig. 4.**
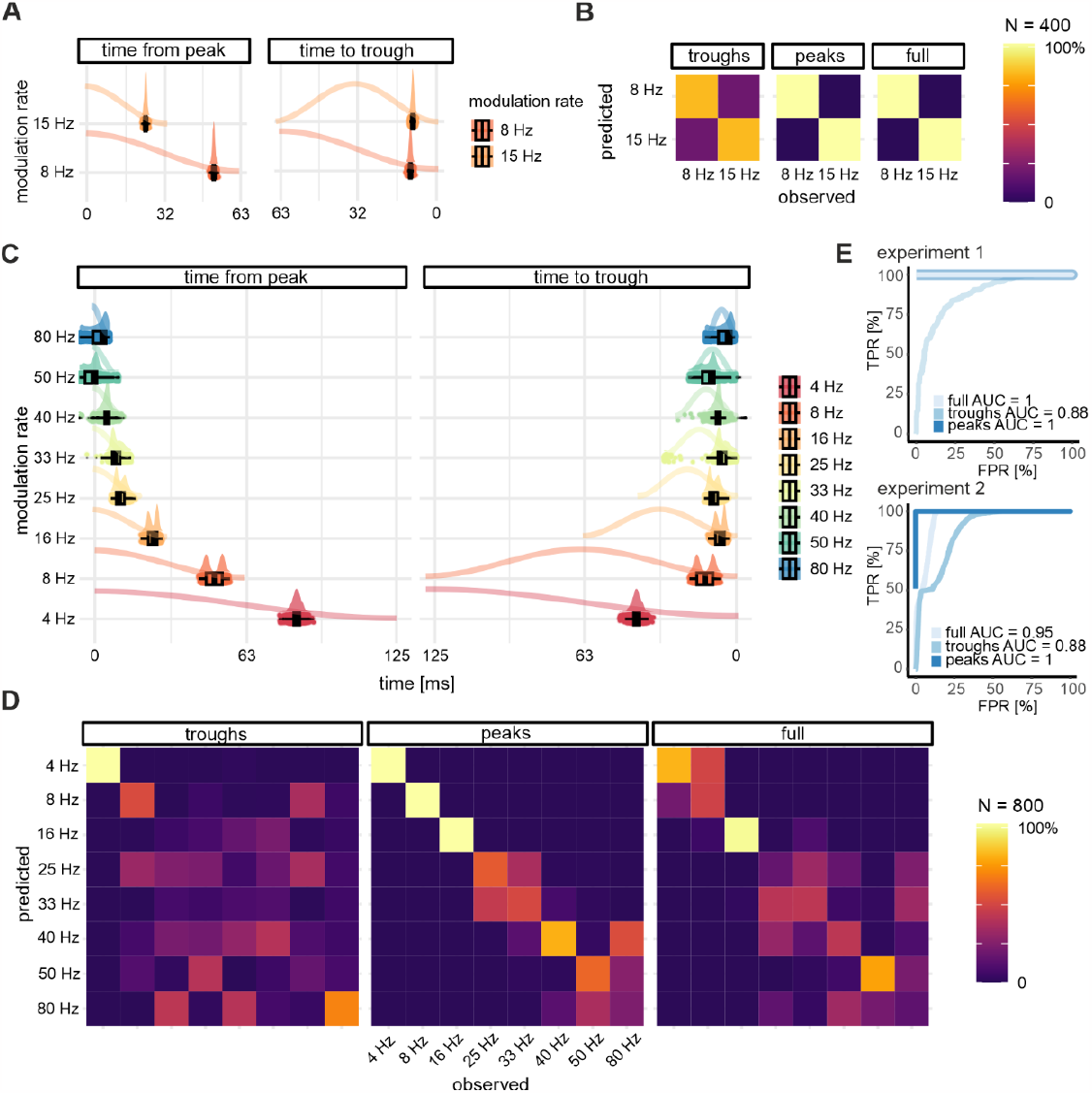
Mean call onset timings measured relative to the amplitude trough and peak show distinct patterns. **(A)** Bootstrapped angular means (in radians) from the broadband masking condition in *experiment 1* expressed as time (in ms) relative to the amplitude peak (left) and relative to the terminal trough (right). Mean call onsets occur in a scaled manner relative to the amplitude peak, but roughly concurrently relative to the terminal trough. Boxplots indicate median, 1^st^ and 3^rd^ quartiles. Whiskers indicate 1.5x the inter-quartile range from box edges. Small dots behind boxplots are raw data values. Half-violins above boxplots are indicate density distributions. Colored traces schematically represent the amplitude envelope. (**B)** Confusion matrices for predicted vs. observed modulation rate classes from three linear discriminant classifiers run on data in **(A)** from *experiment 1*: a “troughs” model using only time-to-trough, a “peaks” models using only time-from-peak, or both. **(C)** Bootstrapped angular means from steady-state and random masking conditions in *experiment 2*. Indications are the same as in **(A). (D**) Same as **(B**), but for data in **(C)** from *experiment 2*. **(E)** ROC curves showing the trade-off between false-positive and true-positive rate for classification by each model.

To adjudicate between these two possibilities, we used a linear classifier to predict modulation rate classes for mean call onsets, using time-from-peak and time-to-trough values as predictors. We ran three models to evaluate the classification performance for each measure separately and together (see Methods): the full model featured both time-to-trough and time-from-peak predictors, the “troughs model” included only the former, and the “peaks model” included only the latter.

Classification performance of an unseen test set was lowest for troughs models for both experiments (Fig. 4B-E, Table 1). The peaks model, using timing relative to amplitude peaks provided perfect (*experiment 1*) or very good classification (*experiment 2*). Finally, a model that used both measures provided perfect performance when the classification task was binary (8 or 15 Hz, *experiment 1*), but performed worse than the peaks model when the task required distinguishing multiple classes (*experiment 2*, Fig. 4B-E, Table 1). A comparison of F1 scores (the geometric mean of precision and recall) confirmed that peaks models performed as well (*experiment 1*) or better (*experiment 2, p* < 0.001) than other models. All models provided significant classification above chance level (*p* < 0.001).

**Table 1.** Classification performance for models predicting modulate rate classes from call onset timing. For experiment 1, F1 scores (geometric mean of precision and recall) are computed based on 8 Hz class being the “positive class.” For experiment 2, the multi-class F1 score is the average of F1 scores for all modulation rate classes.

|  | Model | Accuracy (95% CI) | [Mean] F1 |
| --- | --- | --- | --- |
| **Experiment 1** | **Full model** | 1.0 (0.995, 1) | 1.0 |
|  | **Troughs model** | 0.815 (0.786, 0.841) | 0.806 |
|  | **Peaks model** | 1.0 (0.995, 1) | 1.0 |
| **Experiment 2** | **Full model** | 0.517 (0.504, 0.529) | 0.499 |
|  | **Troughs model** | 0.386 (0.374, 0.398) | 0.363 |
|  | **Peaks model** | 0.709 (0.698, 0.721) | 0.709 |

Together, these results demonstrate that mean call onset timings relative to the terminal amplitude trough were much more similar across modulation rates than timings relative to amplitude peaks, which scaled with cycle length. This difference was most evident for slower modulation rates, for which significant call timing changes were observed. This provides evidence that bats may have calibrated the shift in their vocalization timings by aiming to vocalize at or near the end of the amplitude modulation, when the noise level would have been at a minimum. Given the short duration of most detected calls (above), vocalizations would likely not have extended into the following cycle (Table S17).

Finally, although the masking noise precluded the detection and analysis of returning echoes, we estimate that echoes from echolocation pulses would have arrived at the emitting bat with a delay of 3-7 ms. As median call timings from the terminal trough were on the order of ∽5-40 ms, in conditions where significant vocal timing adjustments were observed, echoes may have incurred even less interference than echolocation pulses.

## Discussion

We found that *Carollia perspicillata* bats adapt to background noise by dynamically adjusting their vocalization timing. Our study makes bats one of several mammals thus far observed to spontaneously adjust to jamming noise by exploiting temporal parameters of the acoustic landscape, alongside marmosets ^23^, cotton-top tamarins ^24^, and dolphins ^25^. This capacity has previously also been found in weakly electric fish ^26^, some songbirds ^27^ and frogs ^28^ as well as numerous insects ^1^, lending evidence to the notion that this ability may be an old and phylogenetically conserved capacity. This adaptation was most evident for modulation rates between 4 and 16 Hz, whether the temporal pattern of amplitude fluctuations was predicable or not. These results invite two interesting inferences.

First, vocal timing in bats is plastic but may be constrained by an intrinsic rate of vocalization, such as the ∽11 Hz rate we observed from our colony. This inference is supported by a previous study which demonstrated that the auditory cortex of bats exhibits phase-locked spiking activity in response to amplitude modulated tones, but only up to ∽22 Hz ^29^. In addition, while our study investigated vocal production behavior while bats were largely stationary, it has long been known that bat vocalizations are temporally linked with respiration, which is in turn coupledwith the wingbeat during flight ^30,31^. Thus, both neural and metabolic constraints may play a role in limiting the range of vocal timing adaptations. However, given that bats of this species are expert echolocators, and Phyllostomid bats have previously been successfully trained to modify social vocalizations ^32^, adaptation to faster rates may be possible under an operant conditioning paradigm. This hypothesis is consistent with our findings which showed diffuse call onset clustering patterns for rates of 25 Hz and above, indicating a gradual roll-off of temporal tracking, rather than a hard cutoff.

Second, the mechanism underlying short-term vocal plasticity is sufficiently fast and flexible to permit adaptation to a range of temporal rates without the need for strict predictability. This is a critical feature, as the natural environment presents numerous acoustic hurdles characterized by erratic temporal patterns, making it highly adaptive to be able to calibrate calling behavior to moment-to-moment fluctuations in amplitude level. In the wild, background noise may also be continuous over long periods of time. While a few studies on bat vocal adaptation have observed phasic vocal responses to playback noise ^33^, or changes in call interval ^34^ and pulse emission timing ^10,11^, these have mainly employed pulsatile or discrete stimuli, rather than continuous playback, to probe vocal production behavior.

An open question raised by our findings regards whether bats achieve jamming avoidance primarily by continuous, active online monitoring of the acoustic landscape, or whether they switch as needed between such a strategy and reference to an internal prediction model of the auditory scene. While an active online monitoring strategy would adequately explain our results, bats have recently been shown to build and act upon predictions of auditory targets ^35^. Future studies are needed to discern whether the bat brain switches between minimally costly approaches to the problem in a context-dependent manner.

Overall, the behavior we observed is consistent with the notion that bats vocalize like metabolically efficient signal optimizers: First, the vocal timing adaptation we observed is employed in a “lazy” manner, i.e. only if the making noise necessitates temporal shifting. It is important to note that masking noise may incur a change in behavior through a combination of direct and indirect interference effects. The spectral frequencies masked by the high-freq masker targeted mainly echolocation pulses, which we estimated to be the majority of detected vocalizations. The fact that we only observed significant call timing changes with the broadband masking noise suggests that direct interference with signal frequencies is not sufficient to incur this behavioral change. This result is analogous to the manner in which Lombard effects increase with more broadband masking noises, even when additional carrier frequencies do not overlap with vocalizations ^4^.

Second, consistent with previous studies, we found that overall rates of calling dropped during playback of masking noise, though this effect was strongly modulated by acoustic context, such that bats appear to evade acoustic jamming by learning global statistical patterns in ambient noise and then locally exploiting the windows of opportunity afforded by slower amplitude fluctuations.

Finally, we found that while mean call onset times scaled with modulation period, call timing with respect to the end of the cycle was more similar between different temporal rates. providing evidence that when bats perform timing adjustments, they do so by aligning call timings to the amplitude trough, where both calls and returning echoes would incur the least amount of interference.

Beyond its importance for maintaining signal quality, vocal flexibility in the temporal domain is a critical prerequisite for complex social communication, as it allows for the ability to respond to conspecific signals of arbitrary length and complexity ^36^. Previous investigations into the functional and anatomical basis of vocal control provide evidence for common or overlapping pathways supporting the production of both vocalization types at the level of the cerebral cortex (namely, the frontal auditory field) ^19,37^, and a differentiation of the motor pathway in the brainstem ^38,39^. Our study may therefore help elucidate how these bats maintain sensitive temporal dynamics under both vocalization regimes.

In sum, our study demonstrates that the Phyllostomid bat *Carollia perspicillata* has a capacity for vocal flexibility in the temporal domain that is finely responsive to continuous and dynamically changing amplitude fluctuations, enabling this species to optimize calling behavior as needed, by integrating acoustic information at the millisecond timescale.

## Methods

### Animals

72 adult bats (24 female) of the species *Carollia perspicillata* were used in this study. Bats were taken from a breeding colony at the Institute for Cell Biology and Neuroscience at Goethe University Frankfurt in Frankfurt am Main, Germany. We have complied with all relevant ethical regulations for animal use. All experiments were conducted in accordance with the Declaration of Helsinki and local regulations in the state of Hessen. The study received ethical approval under experimental permit FU1126 and FR2007, Regierungspräsidium Darmstadt. Animals had access to food (a mixture of banana pulp, oatmeal and honey) and water *ad libitum* when recordings were not taking place. 48 bats (16 female) were used in experiment 1, while 24 (8 female) were used in experiment 2.

### Stimuli

*Experiment 1:*

Two types of masking noise were generated for this experiment, a broadband white noise (carrier frequencies 10-96 kHz) “broadband masker” and a narrower-band (carrier frequencies 50-96 kHz) “high-frequency masker” (60 seconds each). Each noise segment was then amplitude modulated at 8 and 15 Hz, separately. The carrier frequencies for the broadband white noise were selected in order to spectrally mask the peak frequencies used by *Carollia perspicillata* bats for communication calls and echolocation pulses, and only echolocation pulses, respectively^20,40^.

Amplitude modulation rates were chosen to query temporal rates above and below the peak temporal modulation rate of the colony ‘s spontaneous calling, based on previous analysis of acoustic recordings made in the colony (∽11Hz) (Fig. S1).

*Experiment 2:*

A 90 second segment of the broadband masker noise (carrier frequencies 10-96 kHz) was generated and calibrated to account for the dB roll-off induced by the speaker. The calibration curve used to calibrate the stimuli was computed using a custom Matlab GUI (MathWorks), by playing various pure tones through the speaker which were picked up by a Brüel & Kjær microphone positioned roughly at the location in the experimental chamber where the bats tended to congregate. The broadband masker noise was then used to generate eight masking noises with different modulation rates: 4, 8, 16, 25, 33, 40, 50, and 80 Hz. For each modulation rate, we then generated a 7.5 minute long audio file. The eight 7.5 minute files were then randomly permuted and concatenated together to form a 60 minute acoustic stimulus. We then generated a 15 minute long “random masker” by randomly permuting and concatenating together single amplitude modulation cycles for each rate. This was done four times, and the 15 minute sequences were randomly permuted and concatenated together to form a 60 minute acoustic stimulus. The precise sequence in which stimuli were presented was determined by two randomizations, each of which was presented to two groups of bats.

### Procedure

#### Experiment 1

Audio and video were recorded from each of eight groups of bats (4 males, 2 females in each group) in an anechoic chamber (∽120 x 112 x 78 cm) over five consecutive recording days (Fig. S6). On each day, recordings were first made in three one-hour blocks (“playback conditions”): a silent baseline was followed in the second and third blocks by acoustic playback of the broadband and high-freq masking noise (“masking conditions”). (Presentation order of the two masking noises was counterbalanced across groups). Broadband and high-freq masking noise played to each group of bats was either amplitude modulated at 8 or 15 Hz. Hence, each group only ever heard playback noise modulated at one temporal rate, but with different spectral components.

A computer running Matlab 2021a and Avisoft ultrasound recording software (Avisoft-RECORDER USGH, version 4.3.00) controlled simultaneous audio playback, video acquisition and audio recording. A custom Matlab script played the 60-second audio stimuli (16-bit, 192 kHz sampling rate) 60 times to a directional speaker via a RME Fireface 400 FireWire soundcard and amplifier. Stimuli were played at ∽70 dB SPL (measured as root mean square) volume when measured at a distance of ∽30 cm from the speaker. A webcam with infrared filter removed was placed in the cage with a view to the bats’ roosting corner and illuminated by an infrared LED light. Two trigger channels were used to synchronize audio and video recordings with the start of acoustic playback or, in the silent condition, the start of the recording block: the first sent a TTL pulse to the Avisoft recording device (UltraSoundGate 116Hm), which in turn triggered the recording software to begin acquisition from a condenser microphone (250 kHz sampling rate, Avisoft-Bioacoustics CM16); the second illuminated a red photodiode placed in view of the webcam for aligning video and audio offline.

Given the dimensions of the chamber and the ambient temperature, we computed that sound propagation delays between the speaker, microphone, and the bats were all below 1 ms. Additionally, given that the interior of the chamber measured ∽1m^3^, we estimated that the maximum delay between emitted echolocation pulses and returning echoes would be on the order of ∽3-7 ms.

#### Experiment 2

Procedure was similar to that in experiment 1, with the following exceptions: Four groups of bats were tested, each of which was presented with the same acoustic conditions. Acoustic playback in the second and third recording blocks consisted of the steady-state and random masker noise. Precise presentation order and randomizations were counterbalanced between groups. A custom Matlab script controlled simultaneous audio playback, video acquisition and audio recording. 60 minute audio stimuli were played to the speakers.

### Data Analysis

#### Experiment 1

First, any silent periods preceding or following the onset and offset of the masking noise were manually removed (except for groups 1 & 2 in the 8 Hz context, see below). Raw audio files (60 minutes long) were split into segments of 7.5 minutes in duration (all groups except groups 1 & 2 from the 8 Hz condition, for which the raw audio files were 1 minute in duration).

For groups 1 & 2 in the 8 Hz context only: raw data files were saved as 60 second long audio files and featured a brief silence (∽250 ms) at the end of each file, corresponding to the delay caused by the program re-initializing for the next stimulus presentation. These brief silences were trimmed by cross-correlating the amplitude envelope at the end of each file with the amplitude envelope of a recording of the auditory stimulus in the experimental booth without any animals present (“envelope cross-correlation”). Trimmed audio files were visually checked to make sure the end of the file corresponded with the trough of the last amplitude modulation cycle in the file. For files recorded in the silent condition, the final 250 ms of each file was trimmed. For all other groups in this experiment, raw data files were 60 minute long audio files and featured a brief, silent pre- and post-trigger period (∽2 and 0.75 s, respectively). These brief silences were trimmed via envelope cross-correlation. For files recorded in the silent condition, the first 2 and final 0.75 seconds were removed. Files were visually checked and manually edited where the envelope cross-correlation failed to adequately remove artifactual silences.

Vocalization events were detected using *Deep Audio Segmenter* (DAS, v0.28.3) ^41^, a deep neural network developed for the annotation of acoustic signals, and Python (v3.8.3). First, a subset of the dataset was manually annotated. Next, training and test datasets were created from these annotations for the silent and masking conditions, separately. We trained several DAS models using different hyperparameters until we achieved satisfactory prediction and/or a plateau in model improvement. Performance was calculated as the F1 score, the geometric mean of precision and recall. Prediction parameters the same for all runs: 1 ms minimum event duration and 0.9 ms minimum time between event boundaries. Precision, recall, F1 scores, and temporal errors for call onset detection were calculated based on a tolerance of 1.5 ms. Call offsets were detected and used to estimate call durations for the purpose of gaining a broad impression of the proportion of echolocation to communication calls, but otherwise not analyzed, since offsets in our dataset were not very well-defined (i.e. calls frequently featured a decay rather than a sharp offset, or appeared “smeared” due to the appearance of the echo on the recording following the echolocation pulse). Hyperparameters and model performance measures are reported in Tables 2 & 3, respectively.

**Table 2.** Hyperparameters for final models.

|  | Condition | Chunk<br>[samples] | STFT<br>downsample | TCN stacks | Kernel size<br>[samples] | Kernel |
| --- | --- | --- | --- | --- | --- | --- |
| Experiment 1 | Silence | 8192 | 16x | 2 | 16 | 32 |
|  | Masking | 8192 | 16x | 4 | 16 | 32 |
| Experiment 2 | Silence | 8192 | 16x | 2 | 32 | 32 |
|  | Masking | 8192 | 16x | 2 | 16 | 32 |

**Table 3.** Model performance and temporal error in predicting test set.

|  |  | Call Onset Detection |  |  |  |
| --- | --- | --- | --- | --- | --- |
|  |  | Precision | Recall | F1 score | Median<br>temporal error<br>(ms) |
| Experiment 1 | Silence | 0.97 | 0.64 | 0.77 | 0.35 |
|  | Masking |  |  |  |  |
|  | predict 8Hz | 0.94 | 0.65 | 0.75 | 0.37 |
|  | predict 15Hz | 0.96 | 0.45 | 0.61 | 0.38 |
| Experiment 2 | Silence | 0.90 | 0.62 | 0.73 | 0.32 |
|  | Masking | 0.90 | 0.61 | 0.73 | 0.21 |

Finally, we labelled each vocalization event detected in the masking conditions with a value corresponding to the instantaneous phase of the amplitude modulation at the time of call onset. Each audio file was bandpass filtered (10.1-10.5 kHz, 3^rd^ order butterworth filter) to remove acoustic artifacts. The Hilbert envelope of the filtered audio was then downsampled by a factor of 10 and passed through a temporal bandpass filter (modulation rate ±1 Hz, 2^nd^ order butterworth filter) to preserve the amplitude modulation signal while removing other acoustic features. The signal was then demeaned and zero-padded at both ends (20 samples). Next, we detected the troughs of the amplitude modulation signal and used these to reconstruct a phase model (0:2pi) of the amplitude modulation envelope, with each trough as the beginning of the next cycle. Time differences between detected troughs were used to estimate the temporal accuracy of the instantaneous phase model (Fig. S7). Detected vocalizations were finally tagged with the corresponding phase value at call onset. For audio files from the silent condition, calls were tagged according to a simulated instantaneous phase signal modelled as a cosine aligned to the start of the file. This cosine model featured the same modulation rate as the corresponding playback conditions.

#### Experiment 2

First, any silent periods preceding or following the onset and offset of the masking noise were manually removed. For audio files from the silent condition, the first and final 2.2 seconds (corresponding to the pre- and post-trigger period) was removed. Raw audio files from the silent and steady-state conditions were then split into segments of 7.5 minutes in duration, in the latter case corresponding to the playback duration of each individual modulation rate. Files from the random conditions were split into segments of 15 minutes, corresponding to the duration of pseudo-random blocks of modulation rate sequences.

The same procedure was used as in experiment 1 to detect the vocalization events. Model hyperparameters and performance measures are reported in Tables 2 & 3, respectively. To ensure that the model was not biased towards detecting (or failing to detect) vocalizations at particular phases of the amplitude envelope, we computed the instantaneous phase of a subset of predicted call events in the test set (from recordings during the random masker playback, which included samples from all modulation cycles), and grouped them by whether DAS detected a true positive, false positive, or false negative. We found no prominent bias in the detection of call events at any particular phase (Fig. S8).

The same procedure was used as in experiment 1 to label vocalization events with the instantaneous amplitude phase for call detected in the silent and stead-state conditions. For calls detected in the random condition: Each audio file was bandpass filtered (15-60 kHz, 3^rd^ order butterworth filter) to remove acoustic artifacts. The Hilbert envelope of the filtered audio was then downsampled by a factor of 10 and passed through a temporal lowpass filter (70 Hz, 2^nd^ order butterworth filter) to preserve principally the amplitude modulation signal. The signal was then smoothed with a 12-point moving average filter. The sequence of amplitude modulation cycles that comprised the stimulus in each audio file was then used to construct a cosine phase model, which was cross-correlated with the derived modulation signal to obtain an amplitude envelope fit to the recorded audio file. This signal was then demeaned and zero-padded at the ends (20 samples). Next, we detected the troughs of the amplitude modulation signal and used these to reconstruct a phase model (0:2pi) of the amplitude modulation envelope, with each trough as the beginning of the next cycle. Time differences between detected troughs were used to estimate the temporal accuracy of the instantaneous phase model (Fig. S7). Detected vocalizations were finally tagged with the corresponding phase value at call onset. For the random condition, data from all modulation cycles of the same temporal rate were pooled together.

### Statistics and Reproducibility

All statistical analyses were carried out in R (v4.2.1) and RStudio (v2022.7.2.576).

To determine if the presence of amplitude modulated noise affected the timing of emitted calls, we compared the density distribution of call onsets within the real (and in the case of the silent control, simulated) modulation cycle for each playback condition and modulation rate. To describe the distributions in each condition, we computed a battery of circular statistics. These metrics take into consideration the cyclical or circular nature of the data, namely, that values at opposite ends of the linear scale (for phase: 0 and 2π, or for values measured in time: e.g. 0 and 125 ms) represent the same moment in time.

Unimodal circular distributions may be described by treating data points as unit vectors and then computing the direction and length of their resultant vector. Summing these unit vectors gives a resultant vector whose direction is equal to the circular mean,

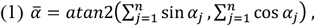

while the length of this vector, given by

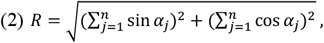

describes how concentrated the data is along the angle given by 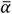. Thus, if provides a measure of *concentration*, then the angular *dispersion* may be defined as *n*−*R* . If is equal to or nearly equal to 0, this indicates that the data are spread evenly over the circumference of the unit circle, and no “preferred direction” exists ^42^. Circular data with a uniform probability density may be called a uniform circular distribution, whose statistical significance can be computed using the Rayleigh test. Meanwhile, the circular normal distribution, called the von Mises distribution, described by a circular mean, μ, and a concentration, *κ*, parameter, can also be fit to the data using maximum likelihood estimation.

We used Mardia-Watson-Wheeler non-parametric tests to test for overall differences in the circular distribution of call onsets between playback conditions within each modulate rate. To simultaneously test for differences in the angular means and angular dispersions, we computed Rao ‘s test of homogeneity. To determine which playback conditions varied significantly from each other on either measure, we computed post-hoc Rao ‘s tests on pairs of conditions where omnibus Rao ‘s tests were significant for either means or dispersions.

These tests were carried out on the entire dataset despite differences in sample size between comparison groups, since the smallest group across both experiments had a sample size of over 5,000 and frequentist circular statistics are only sensitive to sample size at very small Ns ^43^.

To determine whether masking noise influenced the rate of calling, we modelled the number of observed calls in each experimental block (group x recording day x playback condition) using a negative binomial regression using playback condition as predictor, for each modulation rate separately, as follows:

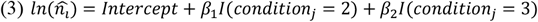

Where *n* is the number of observed call events, *I* is the predictor variable of playback condition with two levels *j* as well as an *intercept* (silence), and *i* is the modulation rate.

In R, models are implemented as follows:

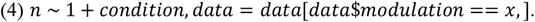

The negative binomial regression was selected for this analysis since a Poisson model with the same formula yielded highly over-dispersed models. Dispersion for all models used was close to 1. A Type II, partial likelihood ratio ANOVA was computed on the negative binomial models to determine significant predictors. Incidence rates and confidence intervals derived from the model were used to estimate the degree to which calling behavior increased or decreased for a given combination of predictors. Post-hoc comparisons evaluated differences in estimated marginal means (predicted calling rates) between pairs of masking conditions. This analysis was repeated for the temporally overlapping calls.

Wherever multiple hypothesis tests were carried out, p-values were adjusted for multiple comparisons by Bonferroni correction. For all hypothesis tests, an alpha level of 0.05 was used.

For the linear discriminant classification analysis, two measures of call onset timing were first computed from bootstrapped angular means as follows:

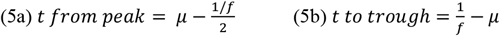

where *μ* is the angular mean computed from an MLE von Mises distribution and *f* is the modulation rate of the current cycle. Thus, time-from-peak values were positive if call onsets occurred on average after the modulation peak in the latter half of the cycle, and negative if call onsets occurred before the peak. Time-to-trough values give the time remaining between average call onset and the final moment in the cycle, the terminal trough. Data from the broadband masking conditions only (experiment 1: broadband masker, experiment 2: steady-state and random masker) was then divided into a training and validation set with 0.6:0.4 split, stratified on modulation rate contexts.

Next, three models were fed the centered (predictor average subtracted from each value) and scaled (predictor values divided by predictor standard deviation) training data and used to determine modulate rate classes predicted by either or both measures, as follows:

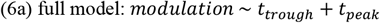

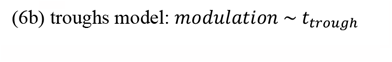

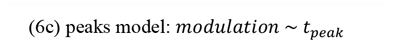

Models (using the *lda* algorithm implemented in the *caret* package) were trained using 10-fold cross-validation (repeated 10 times) and predictors were centered and scaled. Each model was then used to predict modulation rate classes for the validation set. Confusion matrices for observed versus predicted classes from the validation data is shown in Figure 4.

We then computed ROC curves and AUC values for each model from each experiment.

### Manipulated Variables

In experiment 1, one half of the bat groups (4 of 8) were housed together starting on the first recording day and subsequently only for the duration of the experiment (5 days). The other four groups were housed together for a seven-day “familiarization period” prior to the first recording day (12 days in total). An early hypothesis was that groups that did not have the familiarization period may vocalize more, since at the time of data collection all bats that were not part of the experiment were housed separately by sex. Thus, a mixed-sex group could lead to an unusually high level of vocal activity. No clear difference between these two groups emerged based on preliminary results from experiment 1. All analyses were done without respect to this grouping variable.

### Data Exclusion for Experiment 1

For groups 1 & 2 in the 8 Hz context, some recording blocks had buffer issues which caused improper logging of data. Audio files for these blocks were visually checked and sections with corrupted data were removed from the corresponding file if the error was minor (i.e. < 1 second long, or < 3 times per file). If errors were more extensive, the file was removed from analysis. Altogether, approx. 15 minutes of data was removed from the raw data for these two groups combined. For all remaining groups, the first 15 hours of recordings were visually checked for buffer issues. As only a few such occurrences were found, we did not proceed with the visual check.

Original raw data for this experiment amounted to 120 hours of audio recording (approx. 105 hours after data exclusion).

## Acknowledgements

We thank Francisco García-Rosales for advice on circular statistics. We also thank Jonathan Benichov and Luciana López-Jury for helpful discussions. This research was supported by a German Research Foundation grant (DFG Project #275755787 and 428645493) to J.H. and a Main-campus-doctus PhD scholarship (Stiftung Polytechnische Gesellschaft) to A.K. The bat silhouette graphic displayed in Fig. 1a was created by and reproduced here with the permission of Luciana López-Jury.

## Author Contributions

Conceptualization: AK, DP, JH

Methodology: AK, JC, JH

Software: AK, JC

Investigation: AK

Formal Analysis: AK

Visualization: AK

Funding acquisition: AK, JH

Supervision: DP, MK, JH

Writing – original draft: AK

Writing – review & editing: AK, JC, DP, MK, JH

## Data availability

All data used for analysis and figures presented can be obtained from a dedicated repository on [Github](https://github.com/avakiai/colonyjam) (DOI: https://zenodo.org/record/7908545). All other data can be made available from the first corresponding author (AK) upon reasonable request.

## Code availability

All code used for analysis and generation of figures presented can be obtained from a dedicated repository on [Github](https://github.com/avakiai/colonyjam) (DOI: https://zenodo.org/record/7908545).

