## Supplementary Materials for "Flexible control of vocal timing in *Carollia perspicillata* bats enables escape from acoustic interference" (from another version of this preprint: version 2, 2023-10-24)

**The file includes:**

Figs. S1 to S8

Tables S1 to S17

**Supplementary Figures**

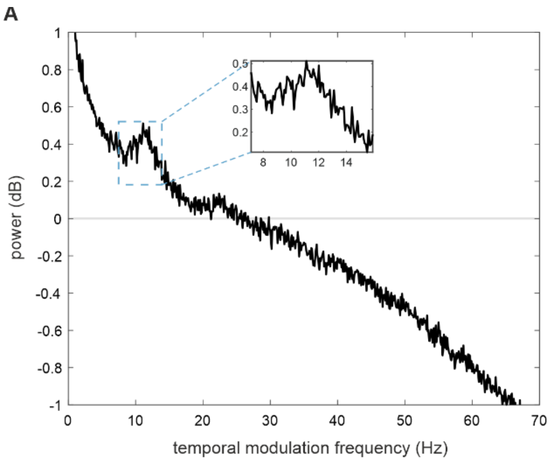


**Fig. S1**.

**Temporal modulation rate of spontaneous vocalizations in C. perspicillata colony is ~ 11 Hz.** (**A)** Audio recordings made in our colony of *Carollia* bats (160 minutes total) were highpass filtered (10 kHz) before amplitude envelopes were extracted via the secant method (interpolation over every 5,000 points). The power spectrum density (PSD) of the envelope was computed for each audio segment (10 s each) and averaged across segments. A "control" PSD was computed by randomly permuting sections of the amplitude envelopes in each segment and averaging the permuted-segment PSDs. The "control" PSD was subtracted from original average PSD to produce a power spectrum normalized for ambient noise/power in the signal. The peak of this power spectrum was at ~ 11 Hz (inset), indicating the temporal modulation rate of spontaneous calling in the colony.

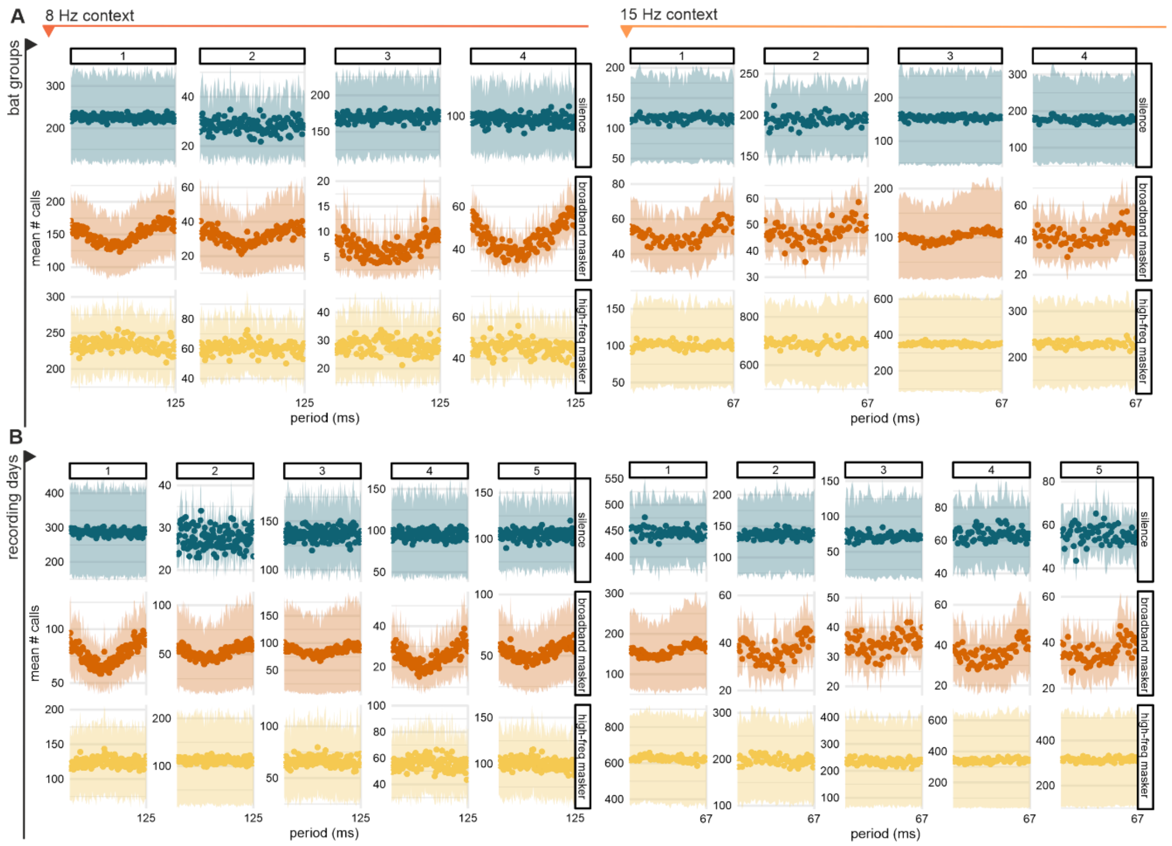


**Fig. S2.**

**Experiment 1: Anti-phase calling is present in all groups and all recording days.**

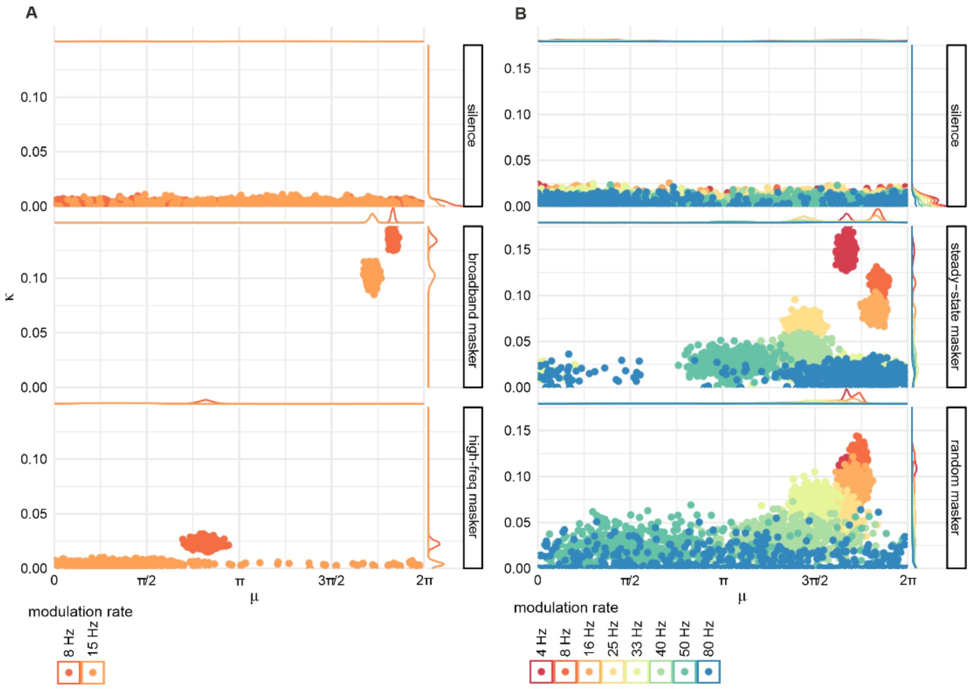


**Fig. S3.**

**Bootstrapped von Mises parameters mean (μ) and concentration (κ) in the cartesian plane. (A)** Parameters from experiment 1 data. **(B)** Parameters from experiment 2 data.

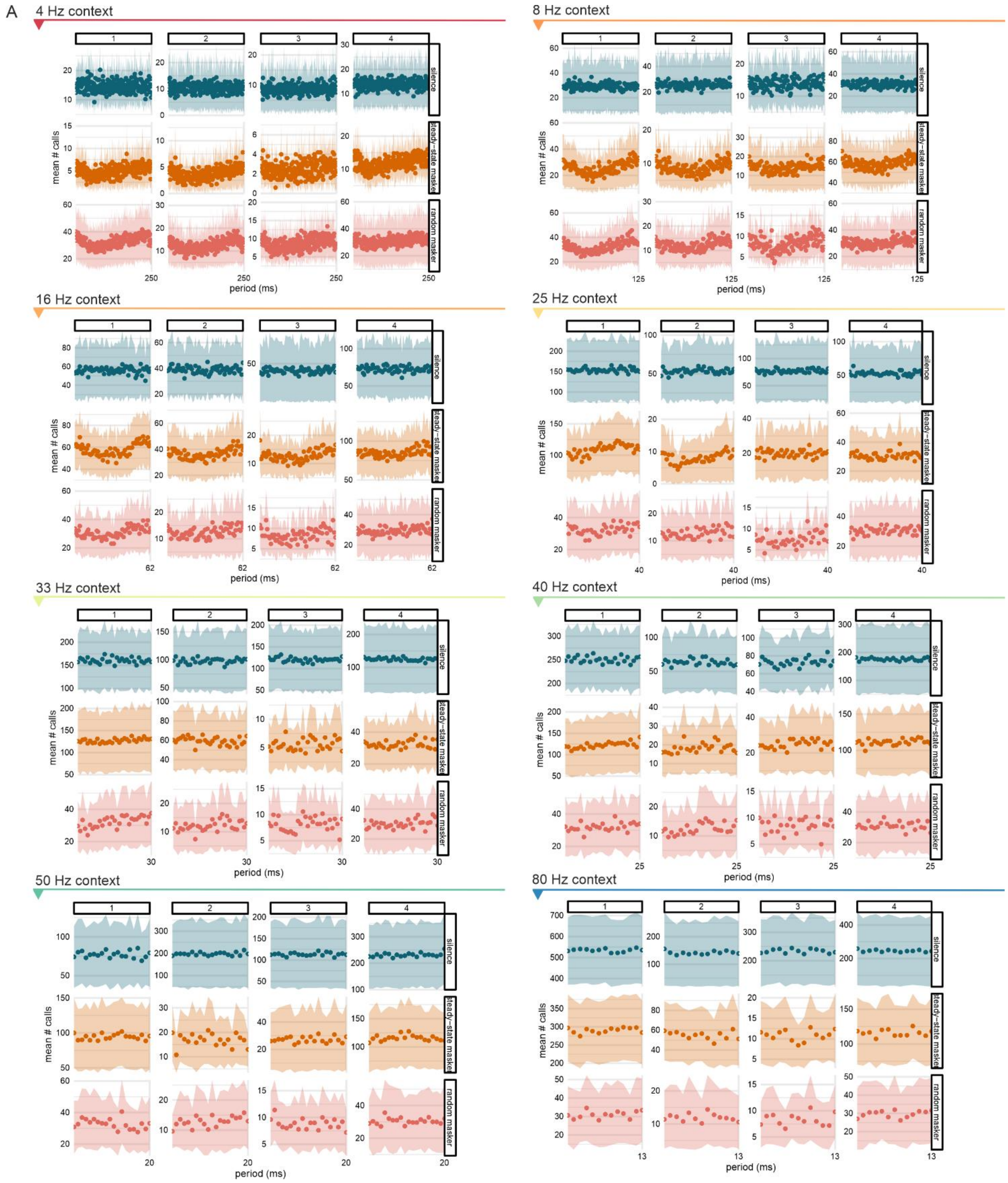


**Fig. S4.**

**Experiment 2: Anti-phase calling is present in all groups for modulation rate conditions 4, 8 and 16. 25 Hz context reveals a slight modulation in calling pattern for some groups between the two masking conditions.**

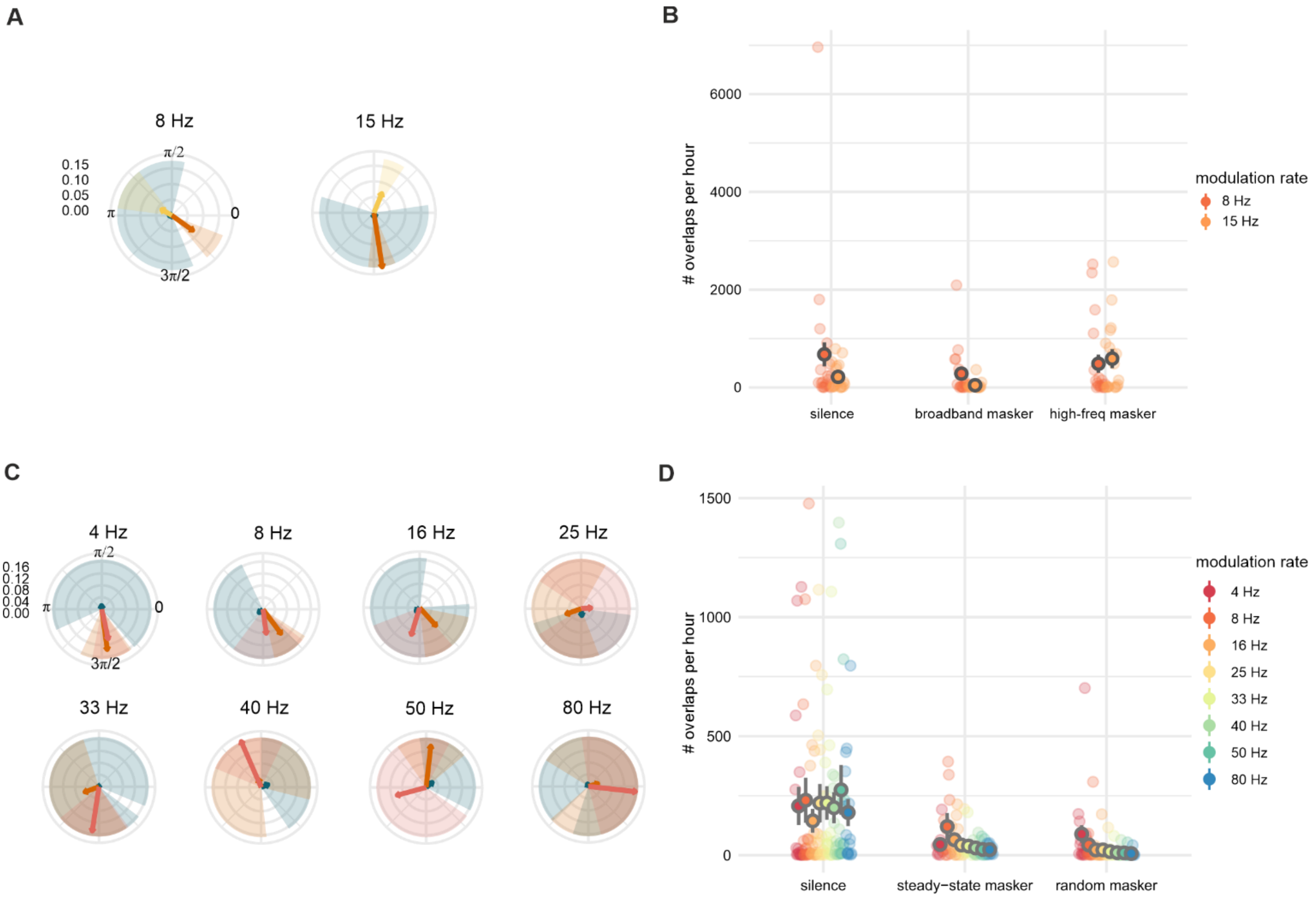


**Fig. S5.**

**Temporal overlap between calls is reduced in the presence of noise. (A)** Angular vectors indicate angular mean and concentration for overlapping calls. Shaded segments indicate 95% confidence intervals for angular means. Significant clustering is seen in the 8Hz broadband and 15Hz high-freq masking condition. **(B)** For experiment 1, estimated mean number of overlapping calls (darkly colored dots with gray outline) observed per hour are reduced in broadband masking noise compared to baseline. Lightly colored dots indicate number of overlaps observed per hour for each group for each recording day. Gray vertical lines indicate SE of model fit for predicted means. **(C)** For experiment 2, overlapping calls are significantly clustered only at 8 and 16Hz in the steady-state condition. (**D**) For experiment 2, estimated mean overlapping calls (darkly colored dots with gray outline) dropped significantly between silent baseline and masking conditions for all modulation rates except 4 Hz. Lightly colored dots indicate number of overlaps observed per hour for each group for each recording day. Gray vertical lines indicate SE of model fit for predicted means.

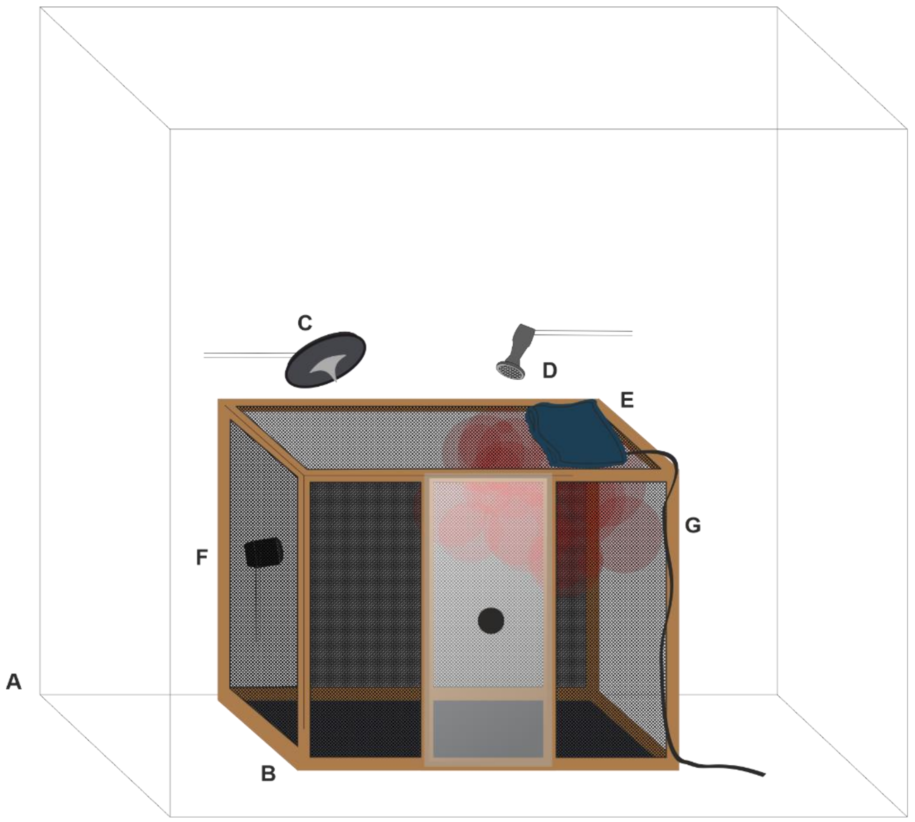


**Fig. S6.**

**Schematic of experimental setup. (a)** Recordings were performed inside a ~120 x 112 x 78 cm (w x h x d, interior measurements), sound-isolated chamber. Bats were placed inside an inner box **(b)** measuring 60 x 47 x 30 cm, where they could freely flit and socialize. A directional speaker **(c)** and Avisoft microphone **(d)** were placed outside the box. A heating pad **(e)** provided warmth and resulted in the bats consistently roosting in the upper right corner of the box. Pink circles **(g)** provide a coarse heatmap (estimated from visual inspection) of the locations where the bats roosted the most. Video was recorded using a webcam placed inside the inner box **(f)** and used to confirm the location of the bats inside the box. Bats most frequently rooted ~30 cm from the speaker and ~15 cm from the microphone. The speaker and microphone were placed approximately 8 cm from each other and 10 cm above the top of the inner box.

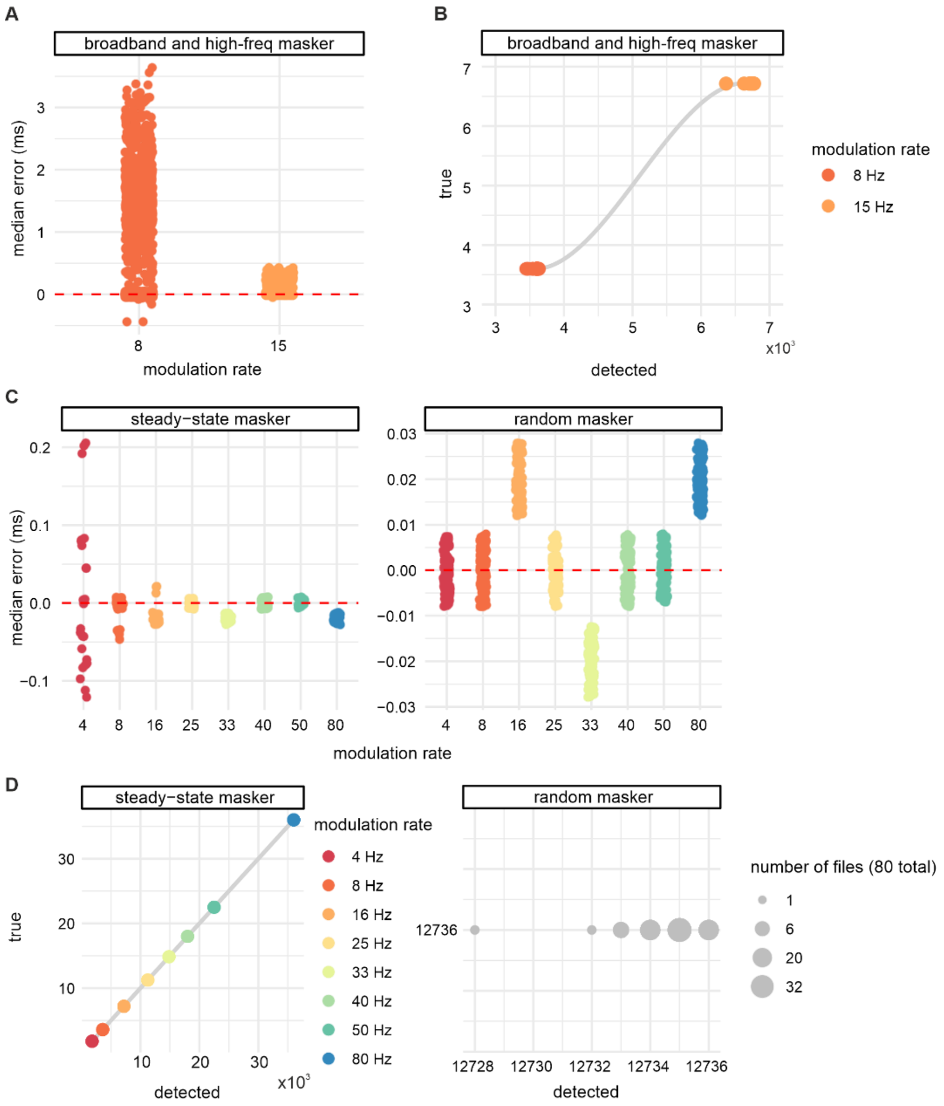


**Fig. S7.**

**Temporal and cycle-wise accuracy of instantaneous phase models. (A)** For experiment 1, the median deviation error between expected modulation periods (1/*f*) and derived periods in the instantaneous phase model for all cycles across the entire data set. Errors were mostly between 0 and 3 ms for both modulation rate contexts. **(B)** For experiment 1, the number of true cycles in each one-hour recording block vs. the number of detected cycles in the instantaneous phase model. On average, 5 more cycles were detected in recording blocks in the 8Hz context, and 19 fewer cycles were detected in the 15Hz contexts. **(C)** For experiment 2, median deviation error between expected and derived periods for each modulation rate were less than ±0.3 ms for all rates in both masking conditions. **(D)** For experiment 2, true vs. detected number of cycles were almost perfectly matched for the steady-state condition. On average, detected cycles were ± 2 cycles from expected values (estimated per 7.5 minute block). For the random condition, true vs. detected cycles were calculated irrespective of modulation rate, but were accurate to 0:-4 cycles per 15 minute block (file).

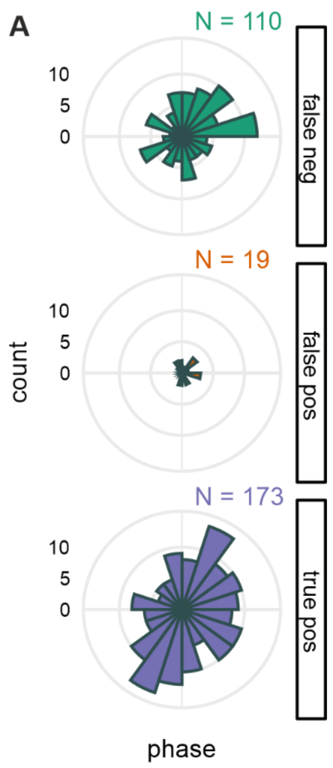


**Fig. S8.**

**DAS-predicted call events by class, as a function of call onset phase.** Phase tagging of detected call events from a representative test set (experiment 2: random masker) showed that DAS is not strongly biased to detect calls at particular phases in the amplitude modulation cycle.

### Supplementary Tables

**Table S1.**

**Estimated proportion of echolocation to communication calls.** A total of 2,673 calls were classified as echolocation or communication calls by visual inspection of spectrograms from all four masking conditions used in both experiments (spectrogram segments sampled from randomly selected groups and files). A weighed average of the proportion of echolocation pulses to communication calls (weighted by the total duration of audio data inspected for each condition) was then computed.

| | Playback Condition | | | |
|---|---|---|---|---|
| | **Silence** | **Broadband masker/Steady-state masker** | **High-freq masker** | **Random masker** |
| Total duration scanned (s) | 180 | 240 | 90 | 90 |
| Condition-wise mean prop. echo | 0.79 | 0.96 | 0.94 | 0.78 |
| Weighted mean prop. echo | | 0.88 | | |

**Table S2.**

**Experiment 1: Circular statistics for call onsets with respect to the amplitude modulation cycle**

| | | Circular Statistics | | | | | | | | | | | |
|---|---|---|---|---|---|---|---|---|---|---|---|---|---|
| | | **Summary Statistics** | | | | | **Rayleigh's Test** | | | **MLE von Mises** | | | |
| **modulation** | **condition** | **n** | $\bar{\theta}$ | $\bar{r}$ | **Vm** | **sd** | **R** | $p$ | $P_{Bonferroni}$ | $\mu_{MLE}$ | **$CI_{lo}$** | **$CI_{hi}$** | $\kappa_{MLE}$ |
| 8Hz | silence | 325,333 | 0.738 | 0.001 | 0.999 | 3.722 | 0.001 | 0.279 | 1 | 0.738 | 4.534 | 4.327 | 0.002 |
| 8Hz | broadband masker | 146,187 | 5.752 | 0.067 | 0.933 | 2.324 | 0.058 | 0.000 | <0.001 | 5.752 | 5.698 | 5.808 | 0.135 |
| 8Hz | high-freq masker | 229,311 | 2.579 | 0.012 | 0.989 | 2.989 | -0.010 | 1.000 | 1 | 2.579 | 2.295 | 2.836 | 0.023 |
| 15Hz | silence | 202,822 | 3.933 | 0.001 | 0.999 | 3.661 | -0.001 | 0.709 | 1 | 3.933 | 1.962 | 0.506 | 0.002 |
| 15Hz | broadband masker | 78,197 | 5.403 | 0.051 | 0.949 | 2.442 | 0.032 | 0.000 | <0.001 | 5.403 | 5.306 | 5.504 | 0.102 |
| 15Hz | high-freq masker | 443,217 | 1.042 | 0.002 | 0.998 | 3.524 | 0.001 | 0.170 | 1 | 1.042 | 6.003 | 2.560 | 0.004 |

N = number of observations. $\bar{\theta}$ = Mean resultant direction. $\bar{r}$ = Mean resultant length. Vm = circular variance. sd = circular standard deviation. R = test statistics for Rayleigh's test of uniformity. p = p value. $P_{Bonferroni}$ = Bonferroni corrected p-values for the family of all tests in the table. $\mu_{MLE}$ = Mean parameter estimated from the maximum likelihood von Mises distribution. $CI_{lo}$, $CI_{hi}$ = Boostrapped upper and lower thresholds for the 95% confidence interval for the MLE mean parameter. $\kappa_{MLE}$ = Concentration parameter estimated from the MLE von Mises distribution.

**Table S3.**

**Experiment 1: Rao's test for differences in angular means or dispersions in distributions of call onsets between playback conditions**

| Rao test for homogeneity of angular means & dispersions within mod rates | | | | | | |
|---|---|---|---|---|---|---|
| **modulation** | **condition** | **test** | **statistic** | **df** | ***p*** | $P_{Bonferroni}$ |
| 8Hz | Omnibus | polar vectors | 0.477 | 2 | 0.788 | 1.000 |
| 8Hz | Omnibus | dispersions | 346.680 | 2 | 0.000 | <0.001 |
| 15Hz | Omnibus | polar vectors | 2.708 | 2 | 0.258 | 0.516 |
| 15Hz | Omnibus | dispersions | 100.345 | 2 | 0.000 | <0.001 |
| 8Hz | silence vs. Broadband masker | polar vectors | 0.422 | 1 | 0.516 | 1.00 |
| 8Hz | silence vs. Broadband masker | dispersions | 332.013 | 1 | 0.000 | <0.001 |
| 8Hz | silence vs. High-freq masker | polar vectors | 0.444 | 1 | 0.505 | 1.00 |
| 8Hz | silence vs. High-freq masker | dispersions | 14.766 | 1 | 0.000 | <0.001 |
| 8Hz | broadband masker vs. High-freq masker | polar vectors | 0.054 | 1 | 0.817 | 1.00 |
| 8Hz | broadband masker vs. High-freq masker | dispersions | 307.346 | 1 | 0.000 | <0.001 |
| 15Hz | silence vs. Broadband masker | polar vectors | 0.739 | 1 | 0.390 | 0.78 |
| 15Hz | silence vs. Broadband masker | dispersions | 100.231 | 1 | 0.000 | <0.001 |
| 15Hz | silence vs. High-freq masker | polar vectors | 0.045 | 1 | 0.832 | 1.00 |
| 15Hz | silence vs. High-freq masker | dispersions | 0.193 | 1 | 0.660 | 1.00 |
| 15Hz | broadband masker vs. High-freq masker | polar vectors | 1.975 | 1 | 0.160 | 0.32 |
| 15Hz | broadband masker vs. High-freq masker | dispersions | 100.029 | 1 | 0.000 | <0.001 |

**Table S4.**

**Experiment 1: Negative binomial model results for calls emitted both modulation rate contexts**

| Observed calls per hour in 8Hz modulated noise | | | |
|---|---|---|---|
| **Predictors** | **Incidence Rate Ratios** | **CI** | ***p*** |
| (Intercept) | 16266.65 | 10048.04 – 28874.24 | <0.001 |
| condition [broadband masker] | 0.45 | 0.21 – 0.95 | 0.034 |
| condition [high-freq masker] | 0.70 | 0.33 – 1.49 | 0.355 |
| Observations | 60 | | |
| $R^2$ Nagelkerke | 0.097 | | |
| Deviance | 72.505 | | |
| AIC | 1239.835 | | |

| Observed calls per hour in 15Hz modulated noise | | | |
|---|---|---|---|
| **Predictors** | **Incidence Rate Ratios** | **CI** | ***p*** |
| (Intercept) | 10674.84 | 6729.32 – 18413.99 | <0.001 |
| condition [broadband masker] | 0.39 | 0.19 – 0.79 | 0.008 |
| condition [high-freq masker] | 2.19 | 1.07 – 4.46 | 0.030 |
| Observations | 57 | | |
| $R^2$ Nagelkerke | 0.395 | | |
| Deviance | 67.547 | | |
| AIC | 1171.104 | | |

Note that the p-values in the table above are not Bonferroni corrected, as they represent p-values for model coefficients.

**Table S5.**

**Experiment 1: Incidence rate ratios for calling derived from negative binomial models**

| Incidence Rate Ratios | | | | |
|---|---|---|---|---|
| **modulation** | **coefficient** | **Estimate** | **2.5 %** | **97.5 %** |
| 8Hz | (Intercept) | 16266.65 | 10048.04 | 28874.24 |
| 8Hz | broadband masker | 0.45 | 0.21 | 0.95 |
| 8Hz | high-freq masker | 0.70 | 0.33 | 1.49 |
| 15Hz | (Intercept) | 10674.84 | 6729.32 | 18413.99 |
| 15Hz | broadband masker | 0.39 | 0.19 | 0.79 |
| 15Hz | high-freq masker | 2.19 | 1.07 | 4.46 |

**Table S6.**

**Experiment 1: ANOVA** Analysis of variance for the predictor of playback condition for each modulation rate model, separately.

| | | Type II Analysis of Deviance | | |
|---|---|---|---|---|
| **model** | **predictor** | **LR Chi-square** | **df** | ***p*** |
| 8Hz | condition | 4.37 | 2 | 0.11 |
| 15Hz | condition | 21.34 | 2 | <0.001 |

**Table S7.**

**Experiment 1: Estimated marginal means of call incidence rates per playback condition**

Estimated Marginal Means

| **modulation** | **contrast** | **IRR** | **se** | **$CI_{lo}$** | **$CI_{hi}$** | **Z ratio** | ***p*** |
|---|---|---|---|---|---|---|---|
| 8Hz | silence / (broadband masker) | 2.22 | 0.84 | 0.90 | 5.50 | 2.12 | 0.10 |
| 8Hz | silence / (high-freq masker) | 1.42 | 0.54 | 0.57 | 3.51 | 0.92 | 1.00 |
| 8Hz | (broadband masker) / (high-freq masker) | 0.64 | 0.24 | 0.26 | 1.58 | -1.19 | 0.70 |
| 15Hz | silence / (broadband masker) | 2.59 | 0.94 | 1.09 | 6.15 | 2.64 | 0.03 |
| 15Hz | silence / (high-freq masker) | 0.46 | 0.16 | 0.19 | 1.08 | -2.17 | 0.09 |
| 15Hz | (broadband masker) / (high-freq masker) | 0.18 | 0.06 | 0.07 | 0.42 | -4.81 | <0.001 |

IRR = Incidence rate ratios.

**Table S8.**

**Experiment 2: Circular statistics for call onsets with respect to the amplitude modulation cycle**

| | | Circular Statistics | | | | | | | | | | | |
|---|---|---|---|---|---|---|---|---|---|---|---|---|---|
| | | **Summary Statistics** | | | | **Rayleigh's Test** | | | | **MLE von Mises** | | | |
| **modulation** | **condition** | **n** | $\bar{\theta}$ | $\bar{r}$ | **Vm** | **sd** | **R** | $p$ | $p_{Bonferroni}$ | $\mu_{MLE}$ | $CI_{lo}$ | $CI_{hi}$ | $\kappa_{MLE}$ |
| 4Hz | silence | 59,635 | 6.203 | 0.003 | 0.997 | 3.388 | 0.003 | 0.134 | 1 | 6.203 | 4.212 | 3.124 | 0.006 |
| 4Hz | steady-state masker | 28,938 | 5.246 | 0.074 | 0.926 | 2.282 | 0.038 | 0.000 | <0.001 | 5.246 | 5.140 | 5.358 | 0.148 |
| 4Hz | random masker | 105,990 | 5.245 | 0.054 | 0.946 | 2.416 | 0.027 | 0.000 | <0.001 | 5.245 | 5.170 | 5.319 | 0.108 |
| 8Hz | silence | 62,203 | 5.480 | 0.002 | 0.998 | 3.462 | 0.002 | 0.270 | 1 | 5.480 | 3.523 | 2.293 | 0.005 |
| 8Hz | steady-state masker | 65,446 | 5.795 | 0.057 | 0.943 | 2.395 | 0.050 | 0.000 | <0.001 | 5.795 | 5.702 | 5.891 | 0.114 |
| 8Hz | random masker | 51,939 | 5.459 | 0.061 | 0.939 | 2.366 | 0.041 | 0.000 | <0.001 | 5.459 | 5.356 | 5.554 | 0.122 |
| 16Hz | silence | 65,769 | 0.858 | 0.003 | 0.997 | 3.376 | 0.002 | 0.213 | 1 | 0.858 | 5.075 | 3.812 | 0.007 |
| 16Hz | steady-state masker | 60,482 | 5.732 | 0.042 | 0.958 | 2.520 | 0.036 | 0.000 | <0.001 | 5.732 | 5.603 | 5.871 | 0.084 |
| 16Hz | random masker | 25,543 | 5.376 | 0.046 | 0.955 | 2.486 | 0.028 | 0.000 | <0.001 | 5.376 | 5.178 | 5.575 | 0.091 |
| 25Hz | silence | 67,403 | 4.819 | 0.003 | 0.997 | 3.373 | 0.000 | 0.448 | 1 | 4.819 | 2.541 | 0.274 | 0.007 |
| 25Hz | steady-state masker | 32,743 | 4.535 | 0.032 | 0.968 | 2.618 | -0.006 | 0.928 | 1 | 4.535 | 4.299 | 4.766 | 0.065 |
| 25Hz | random masker | 16,600 | 5.056 | 0.027 | 0.973 | 2.689 | 0.009 | 0.050 | 1 | 5.056 | 4.653 | 5.478 | 0.054 |
| 33Hz | silence | 78,179 | 0.895 | 0.004 | 0.996 | 3.313 | 0.003 | 0.153 | 1 | 0.895 | 5.646 | 2.632 | 0.008 |
| 33Hz | steady-state masker | 34,843 | 5.359 | 0.008 | 0.992 | 3.093 | 0.005 | 0.092 | 1 | 5.359 | 4.466 | 0.165 | 0.017 |
| 33Hz | random masker | 12,830 | 4.688 | 0.034 | 0.966 | 2.599 | -0.001 | 0.552 | 1 | 4.688 | 4.358 | 5.069 | 0.068 |
| 40Hz | silence | 68,797 | 0.196 | 0.001 | 0.999 | 3.668 | 0.001 | 0.331 | 1 | 0.196 | 4.283 | 2.335 | 0.002 |
| 40Hz | steady-state masker | 34,176 | 4.486 | 0.020 | 0.980 | 2.801 | -0.004 | 0.877 | 1 | 4.486 | 4.063 | 4.874 | 0.040 |
| 40Hz | random masker | 10,531 | 4.241 | 0.015 | 0.985 | 2.891 | -0.007 | 0.844 | 1 | 4.241 | 3.404 | 5.347 | 0.031 |
| 50Hz | silence | 61,956 | 5.052 | 0.002 | 0.998 | 3.564 | 0.001 | 0.419 | 1 | 5.052 | 2.940 | 1.454 | 0.004 |
| 50Hz | steady-state masker | 25,425 | 3.302 | 0.013 | 0.987 | 2.937 | -0.013 | 0.999 | 1 | 3.302 | 2.584 | 4.017 | 0.027 |
| 50Hz | random masker | 8,487 | 1.776 | 0.009 | 0.991 | 3.055 | -0.002 | 0.599 | 1 | 1.776 | 5.797 | 3.788 | 0.019 |
| 80Hz | silence | 75,144 | 0.679 | 0.002 | 0.998 | 3.457 | 0.002 | 0.221 | 1 | 0.679 | 4.565 | 3.322 | 0.005 |
| 80Hz | steady-state masker | 30,675 | 5.427 | 0.007 | 0.993 | 3.166 | 0.004 | 0.139 | 1 | 5.427 | 3.897 | 0.352 | 0.013 |
| 80Hz | random masker | 5,260 | 4.324 | 0.002 | 0.998 | 3.561 | -0.001 | 0.527 | 1 | 4.324 | 1.952 | 2.322 | 0.004 |

**Table S9.**

**Experiment 2: Watson-Wheeler tests for the circular homogeneity of distributions of call onsets**

| Watson-Wheeler test for homogeneity of groups (within modulation rates) | | | |
|---|---|---|---|
| **modulation** | **statistic** | ***p*** | $p_{Bonferroni}$ |
| 4Hz | 281.693 | <0.001 | <0.001 |
| 8Hz | 273.637 | <0.001 | <0.001 |
| 16Hz | 143.040 | <0.001 | <0.001 |
| 25Hz | 45.416 | <0.001 | <0.001 |
| 33Hz | 32.695 | <0.001 | <0.001 |
| 40Hz | 20.606 | <0.001 | <0.001 |
| 50Hz | 8.775 | 0.067 | 0.536 |
| 80Hz | 2.286 | 0.683 | 1.000 |

**Table S10.**

**Experiment 2: Rao's test for differences in angular means or dispersions between playback conditions**
Omnibus tests were carried out for each modulation rate to test if differences existed between the three playback conditions. For all tests with significant differences in either polar vectors or polar dispersions, post-hoc tests were computed on pairs of playback conditions.

Rao test for homogeneity of angular means & dispersions

| modulation | condition | test | statistic | df | *p* | $p_{Bonferroni}$ |
|---|---|---|---|---|---|---|
| 4Hz | Omnibus | polar vectors | 3.110 | 2 | 0.211 | 0.422 |
| 4Hz | Omnibus | dispersions | 232.370 | 2 | <0.001 | <0.001 |
| 8Hz | Omnibus | polar vectors | 18.883 | 2 | <0.001 | <0.001 |
| 8Hz | Omnibus | dispersions | 199.932 | 2 | <0.001 | <0.001 |
| 16Hz | Omnibus | polar vectors | 6.895 | 2 | 0.032 | 0.064 |
| 16Hz | Omnibus | dispersions | 77.400 | 2 | <0.001 | <0.001 |
| 25Hz | Omnibus | polar vectors | 3.853 | 2 | 0.146 | 0.292 |
| 25Hz | Omnibus | dispersions | 22.588 | 2 | <0.001 | <0.001 |
| 33Hz | Omnibus | polar vectors | 1.668 | 2 | 0.434 | 0.868 |
| 33Hz | Omnibus | dispersions | 7.870 | 2 | 0.020 | 0.040 |
| 40Hz | Omnibus | polar vectors | 0.897 | 2 | 0.639 | 1.000 |
| 40Hz | Omnibus | dispersions | 7.848 | 2 | 0.020 | 0.040 |
| 50Hz | Omnibus | polar vectors | 0.106 | 2 | 0.948 | 1.000 |
| 50Hz | Omnibus | dispersions | 2.521 | 2 | 0.284 | 0.568 |
| 80Hz | Omnibus | polar vectors | 0.806 | 2 | 0.668 | 1.000 |
| 80Hz | Omnibus | dispersions | 0.492 | 2 | 0.782 | 1.000 |
| 4Hz | silence vs. steady-state masker | polar vectors | 2.985 | 1 | 0.084 | 0.168 |
| 4Hz | silence vs. steady-state masker | dispersions | 79.850 | 1 | 0.000 | 0.000 |
| 4Hz | silence vs. random masker | polar vectors | 3.087 | 1 | 0.079 | 0.158 |
| 4Hz | silence vs. random masker | dispersions | 153.051 | 1 | 0.000 | 0.000 |
| 4Hz | steady-state masker vs. random masker | polar vectors | 0.000 | 1 | 0.983 | 1.000 |
| 4Hz | steady-state masker vs. random masker | dispersions | 15.231 | 1 | 0.000 | 0.000 |
| 8Hz | silence vs. steady-state masker | polar vectors | 0.046 | 1 | 0.830 | 1.000 |
| 8Hz | silence vs. steady-state masker | dispersions | 104.750 | 1 | 0.000 | 0.000 |
| 8Hz | silence vs. random masker | polar vectors | 0.000 | 1 | 0.985 | 1.000 |
| 8Hz | silence vs. random masker | dispersions | 95.520 | 1 | 0.000 | 0.000 |
| 8Hz | steady-state masker vs. random masker | polar vectors | 18.858 | 1 | 0.000 | 0.000 |
| 8Hz | steady-state masker vs. random masker | dispersions | 0.944 | 1 | 0.331 | 0.662 |
| 8Hz | silence vs. steady-state masker | polar vectors | 0.046 | 1 | 0.830 | 1.000 |
| 8Hz | silence vs. steady-state masker | dispersions | 104.750 | 1 | 0.000 | 0.000 |
| 8Hz | silence vs. random masker | polar vectors | 0.000 | 1 | 0.985 | 1.000 |
| 8Hz | silence vs. random masker | dispersions | 95.520 | 1 | 0.000 | 0.000 |
| 8Hz | steady-state masker vs. random masker | polar vectors | 18.858 | 1 | 0.000 | 0.000 |
| 8Hz | steady-state masker vs. random masker | dispersions | 0.944 | 1 | 0.331 | 0.662 |

**Table S11.**

**Experiment 2: Estimated marginal means of call incidence rates across modulation rates**

| Estimated Marginal Means | | | | | | | |
|---|---|---|---|---|---|---|---|
| **modulation** | **contrast** | **IRR** | **se** | **$CI_{lo}$** | **$CI_{hi}$** | **Z ratio** | ***p*** |
| silence | 4Hz / 8Hz | 0.96 | 0.47 | 0.20 | 4.49 | -0.09 | 1.00 |
| silence | 4Hz / 16Hz | 0.91 | 0.45 | 0.19 | 4.24 | -0.20 | 1.00 |
| silence | 4Hz / 25Hz | 0.88 | 0.44 | 0.19 | 4.14 | -0.25 | 1.00 |
| silence | 4Hz / 33Hz | 0.76 | 0.38 | 0.16 | 3.57 | -0.55 | 1.00 |
| silence | 4Hz / 40Hz | 0.82 | 0.41 | 0.17 | 3.93 | -0.39 | 1.00 |
| silence | 4Hz / 50Hz | 0.96 | 0.48 | 0.21 | 4.50 | -0.08 | 1.00 |
| silence | 4Hz / 80Hz | 0.79 | 0.39 | 0.17 | 3.71 | -0.47 | 1.00 |
| silence | 8Hz / 16Hz | 0.95 | 0.47 | 0.20 | 4.43 | -0.11 | 1.00 |
| silence | 8Hz / 25Hz | 0.92 | 0.46 | 0.20 | 4.32 | -0.16 | 1.00 |
| silence | 8Hz / 33Hz | 0.80 | 0.39 | 0.17 | 3.72 | -0.46 | 1.00 |
| silence | 8Hz / 40Hz | 0.86 | 0.43 | 0.18 | 4.10 | -0.30 | 1.00 |
| silence | 8Hz / 50Hz | 1.00 | 0.50 | 0.22 | 4.70 | 0.01 | 1.00 |
| silence | 8Hz / 80Hz | 0.83 | 0.41 | 0.18 | 3.87 | -0.38 | 1.00 |
| silence | 16Hz / 25Hz | 0.98 | 0.48 | 0.21 | 4.57 | -0.05 | 1.00 |
| silence | 16Hz / 33Hz | 0.84 | 0.42 | 0.18 | 3.94 | -0.35 | 1.00 |
| silence | 16Hz / 40Hz | 0.91 | 0.46 | 0.19 | 4.34 | -0.19 | 1.00 |
| silence | 16Hz / 50Hz | 1.06 | 0.52 | 0.23 | 4.97 | 0.12 | 1.00 |
| silence | 16Hz / 80Hz | 0.88 | 0.43 | 0.19 | 4.10 | -0.27 | 1.00 |
| silence | 25Hz / 33Hz | 0.86 | 0.43 | 0.18 | 4.04 | -0.30 | 1.00 |
| silence | 25Hz / 40Hz | 0.93 | 0.47 | 0.20 | 4.44 | -0.14 | 1.00 |
| silence | 25Hz / 50Hz | 1.09 | 0.54 | 0.23 | 5.09 | 0.17 | 1.00 |
| silence | 25Hz / 80Hz | 0.90 | 0.44 | 0.19 | 4.20 | -0.22 | 1.00 |
| silence | 33Hz / 40Hz | 1.08 | 0.54 | 0.23 | 5.16 | 0.15 | 1.00 |
| silence | 33Hz / 50Hz | 1.26 | 0.62 | 0.27 | 5.90 | 0.47 | 1.00 |
| silence | 33Hz / 80Hz | 1.04 | 0.51 | 0.22 | 4.87 | 0.08 | 1.00 |
| silence | 40Hz / 50Hz | 1.17 | 0.58 | 0.24 | 5.58 | 0.31 | 1.00 |
| silence | 40Hz / 80Hz | 0.96 | 0.48 | 0.20 | 4.60 | -0.07 | 1.00 |
| silence | 50Hz / 80Hz | 0.82 | 0.41 | 0.18 | 3.86 | -0.39 | 1.00 |
| steady-state masker | 4Hz / 8Hz | 0.42 | 0.21 | 0.09 | 2.05 | -1.71 | 1.00 |
| steady-state masker | 4Hz / 16Hz | 0.48 | 0.24 | 0.10 | 2.29 | -1.47 | 1.00 |
| steady-state masker | 4Hz / 25Hz | 0.84 | 0.43 | 0.17 | 4.10 | -0.34 | 1.00 |
| steady-state masker | 4Hz / 33Hz | 0.83 | 0.42 | 0.17 | 3.97 | -0.37 | 1.00 |
| steady-state masker | 4Hz / 40Hz | 0.80 | 0.41 | 0.16 | 3.93 | -0.43 | 1.00 |
| steady-state masker | 4Hz / 50Hz | 1.14 | 0.57 | 0.24 | 5.44 | 0.26 | 1.00 |
| steady-state masker | 4Hz / 80Hz | 0.94 | 0.47 | 0.20 | 4.51 | -0.12 | 1.00 |
| steady-state masker | 8Hz / 16Hz | 1.14 | 0.58 | 0.23 | 5.56 | 0.26 | 1.00 |
| steady-state masker | 8Hz / 25Hz | 2.00 | 1.03 | 0.40 | 9.95 | 1.35 | 1.00 |
| steady-state masker | 8Hz / 33Hz | 1.98 | 1.00 | 0.41 | 9.65 | 1.34 | 1.00 |
| steady-state masker | 8Hz / 40Hz | 1.92 | 0.98 | 0.38 | 9.54 | 1.26 | 1.00 |
| steady-state masker | 8Hz / 50Hz | 2.71 | 1.38 | 0.56 | 13.22 | 1.96 | 1.00 |
| steady-state masker | 8Hz / 80Hz | 2.25 | 1.14 | 0.46 | 10.96 | 1.59 | 1.00 |
| steady-state masker | 16Hz / 25Hz | 1.75 | 0.89 | 0.36 | 8.56 | 1.11 | 1.00 |
| steady-state masker | 16Hz / 33Hz | 1.74 | 0.87 | 0.36 | 8.30 | 1.10 | 1.00 |
| steady-state masker | 16Hz / 40Hz | 1.68 | 0.85 | 0.34 | 8.20 | 1.02 | 1.00 |
| steady-state masker | 16Hz / 50Hz | 2.38 | 1.19 | 0.50 | 11.37 | 1.73 | 1.00 |
| steady-state masker | 16Hz / 80Hz | 1.97 | 0.99 | 0.41 | 9.43 | 1.35 | 1.00 |
| steady-state masker | 25Hz / 33Hz | 0.99 | 0.50 | 0.20 | 4.83 | -0.02 | 1.00 |
| steady-state masker | 25Hz / 40Hz | 0.96 | 0.49 | 0.19 | 4.77 | -0.08 | 1.00 |
| steady-state masker | 25Hz / 50Hz | 1.36 | 0.69 | 0.28 | 6.62 | 0.60 | 1.00 |
| steady-state masker | 25Hz / 80Hz | 1.12 | 0.57 | 0.23 | 5.48 | 0.23 | 1.00 |
| steady-state masker | 33Hz / 40Hz | 0.97 | 0.49 | 0.20 | 4.73 | -0.06 | 1.00 |

Estimated Marginal Means

| **modulation** | **contrast** | **IRR** | **se** | **$CI_{lo}$** | **$CI_{hi}$** | **Z ratio** | ***p*** |
|---|---|---|---|---|---|---|---|
| steady-state masker | 33Hz / 50Hz | 1.37 | 0.69 | 0.29 | 6.55 | 0.63 | 1.00 |
| steady-state masker | 33Hz / 80Hz | 1.14 | 0.57 | 0.24 | 5.43 | 0.25 | 1.00 |
| steady-state masker | 40Hz / 50Hz | 1.42 | 0.72 | 0.29 | 6.91 | 0.68 | 1.00 |
| steady-state masker | 40Hz / 80Hz | 1.17 | 0.60 | 0.24 | 5.72 | 0.31 | 1.00 |
| steady-state masker | 50Hz / 80Hz | 0.83 | 0.42 | 0.17 | 3.96 | -0.38 | 1.00 |
| random masker | 4Hz / 8Hz | 2.04 | 0.88 | 0.53 | 7.82 | 1.66 | 1.00 |
| random masker | 4Hz / 16Hz | 4.15 | 1.78 | 1.08 | 15.89 | 3.31 | 0.03 |
| random masker | 4Hz / 25Hz | 6.38 | 2.74 | 1.67 | 24.46 | 4.31 | <0.001 |
| random masker | 4Hz / 33Hz | 8.26 | 3.55 | 2.16 | 31.65 | 4.91 | <0.001 |
| random masker | 4Hz / 40Hz | 10.06 | 4.33 | 2.63 | 38.56 | 5.37 | <0.001 |
| random masker | 4Hz / 50Hz | 12.49 | 5.37 | 3.26 | 47.85 | 5.87 | <0.001 |
| random masker | 4Hz / 80Hz | 20.15 | 8.67 | 5.26 | 77.22 | 6.98 | <0.001 |
| random masker | 8Hz / 16Hz | 2.03 | 0.87 | 0.53 | 7.79 | 1.65 | 1.00 |
| random masker | 8Hz / 25Hz | 3.13 | 1.34 | 0.82 | 11.98 | 2.65 | 0.22 |
| random masker | 8Hz / 33Hz | 4.05 | 1.74 | 1.06 | 15.51 | 3.25 | 0.03 |
| random masker | 8Hz / 40Hz | 4.93 | 2.12 | 1.29 | 18.90 | 3.71 | 0.01 |
| random masker | 8Hz / 50Hz | 6.12 | 2.63 | 1.60 | 23.45 | 4.21 | <0.001 |
| random masker | 8Hz / 80Hz | 9.87 | 4.25 | 2.58 | 37.84 | 5.32 | <0.001 |
| random masker | 16Hz / 25Hz | 1.54 | 0.66 | 0.40 | 5.89 | 1.00 | 1.00 |
| random masker | 16Hz / 33Hz | 1.99 | 0.86 | 0.52 | 7.63 | 1.60 | 1.00 |
| random masker | 16Hz / 40Hz | 2.43 | 1.04 | 0.63 | 9.29 | 2.06 | 1.00 |
| random masker | 16Hz / 50Hz | 3.01 | 1.29 | 0.78 | 11.53 | 2.56 | 0.29 |
| random masker | 16Hz / 80Hz | 4.86 | 2.09 | 1.27 | 18.61 | 3.67 | 0.01 |
| random masker | 25Hz / 33Hz | 1.29 | 0.56 | 0.34 | 4.96 | 0.60 | 1.00 |
| random masker | 25Hz / 40Hz | 1.58 | 0.68 | 0.41 | 6.04 | 1.06 | 1.00 |
| random masker | 25Hz / 50Hz | 1.96 | 0.84 | 0.51 | 7.50 | 1.56 | 1.00 |
| random masker | 25Hz / 80Hz | 3.16 | 1.36 | 0.82 | 12.10 | 2.67 | 0.21 |
| random masker | 33Hz / 40Hz | 1.22 | 0.52 | 0.32 | 4.67 | 0.46 | 1.00 |
| random masker | 33Hz / 50Hz | 1.51 | 0.65 | 0.39 | 5.79 | 0.96 | 1.00 |
| random masker | 33Hz / 80Hz | 2.44 | 1.05 | 0.64 | 9.35 | 2.07 | 1.00 |
| random masker | 40Hz / 50Hz | 1.24 | 0.53 | 0.32 | 4.76 | 0.50 | 1.00 |
| random masker | 40Hz / 80Hz | 2.00 | 0.86 | 0.52 | 7.68 | 1.61 | 1.00 |
| random masker | 50Hz / 80Hz | 1.61 | 0.69 | 0.42 | 6.19 | 1.11 | 1.00 |

**Table S12.**

**Experiment 2: Estimated marginal means for incidence rates of calling**

Estimated Marginal Means

| modulation | contrast | IRR | se | $CI_{lo}$ | $CI_{hi}$ | Z ratio | *p* |
|---|---|---|---|---|---|---|---|
| 4Hz | silence / (steady-state masker) | 2.06 | 0.95 | 0.69 | 6.20 | 1.57 | 0.35 |
| 4Hz | silence / random masker | 0.56 | 0.26 | 0.19 | 1.69 | -1.25 | 0.63 |
| 4Hz | (steady-state masker) / random masker | 0.27 | 0.13 | 0.09 | 0.82 | -2.82 | 0.01 |
| 8Hz | silence / (steady-state masker) | 0.90 | 0.47 | 0.26 | 3.10 | -0.20 | 1.00 |
| 8Hz | silence / random masker | 1.20 | 0.61 | 0.36 | 4.04 | 0.36 | 1.00 |
| 8Hz | (steady-state masker) / random masker | 1.33 | 0.68 | 0.39 | 4.55 | 0.55 | 1.00 |
| 16Hz | silence / (steady-state masker) | 1.09 | 0.48 | 0.38 | 3.10 | 0.19 | 1.00 |
| 16Hz | silence / random masker | 2.58 | 1.13 | 0.90 | 7.34 | 2.16 | 0.09 |
| 16Hz | (steady-state masker) / random masker | 2.37 | 1.04 | 0.83 | 6.75 | 1.97 | 0.15 |
| 25Hz | silence / (steady-state masker) | 1.96 | 0.99 | 0.58 | 6.54 | 1.33 | 0.55 |
| 25Hz | silence / random masker | 4.06 | 2.02 | 1.23 | 13.36 | 2.82 | 0.01 |
| 25Hz | (steady-state masker) / random masker | 2.08 | 1.05 | 0.62 | 6.94 | 1.45 | 0.44 |
| 33Hz | silence / (steady-state masker) | 2.24 | 1.09 | 0.70 | 7.16 | 1.67 | 0.29 |
| 33Hz | silence / random masker | 6.09 | 2.95 | 1.91 | 19.44 | 3.73 | <0.001 |
| 33Hz | (steady-state masker) / random masker | 2.72 | 1.32 | 0.85 | 8.66 | 2.06 | 0.12 |
| 40Hz | silence / (steady-state masker) | 2.01 | 0.95 | 0.65 | 6.26 | 1.48 | 0.42 |
| 40Hz | silence / random masker | 6.88 | 3.22 | 2.24 | 21.08 | 4.12 | <0.001 |
| 40Hz | (steady-state masker) / random masker | 3.42 | 1.60 | 1.11 | 10.47 | 2.62 | 0.03 |
| 50Hz | silence / (steady-state masker) | 2.44 | 1.22 | 0.74 | 8.04 | 1.79 | 0.22 |
| 50Hz | silence / random masker | 7.30 | 3.64 | 2.21 | 24.08 | 3.99 | <0.001 |
| 50Hz | (steady-state masker) / random masker | 3.00 | 1.49 | 0.91 | 9.88 | 2.20 | 0.08 |
| 80Hz | silence / (steady-state masker) | 2.45 | 1.11 | 0.82 | 7.28 | 1.97 | 0.15 |
| 80Hz | silence / random masker | 14.29 | 6.50 | 4.80 | 42.50 | 5.84 | <0.001 |
| 80Hz | (steady-state masker) / random masker | 5.83 | 2.66 | 1.96 | 17.35 | 3.87 | <0.001 |

**Table S13.**

**Experiment 2: Incidence rate ratios derived from negative binomial models**

| | | Incidence Rate Ratios | | |
|---|---|---|---|---|
| **modulation** | **coefficient** | **Estimate** | **2.5 %** | **97.5 %** |
| 4Hz | (Intercept) | 2981.75 | 1675.19 | 6085.74 |
| 4Hz | steady-state masker | 0.49 | 0.19 | 1.21 |
| 4Hz | random masker | 1.78 | 0.71 | 4.45 |
| 8Hz | (Intercept) | 3110.15 | 1655.22 | 6909.31 |
| 8Hz | steady-state masker | 1.11 | 0.40 | 3.12 |
| 8Hz | random masker | 0.83 | 0.30 | 2.31 |
| 16Hz | (Intercept) | 3288.45 | 1895.48 | 6456.38 |
| 16Hz | steady-state masker | 0.92 | 0.38 | 2.20 |
| 16Hz | random masker | 0.39 | 0.16 | 0.93 |
| 25Hz | (Intercept) | 3370.15 | 1815.43 | 7344.02 |
| 25Hz | steady-state masker | 0.51 | 0.19 | 1.41 |
| 25Hz | random masker | 0.25 | 0.09 | 0.67 |
| 33Hz | (Intercept) | 3908.95 | 2136.58 | 8324.83 |
| 33Hz | steady-state masker | 0.45 | 0.17 | 1.17 |
| 33Hz | random masker | 0.16 | 0.06 | 0.43 |
| 40Hz | (Intercept) | 3620.89 | 2003.11 | 7568.02 |
| 40Hz | steady-state masker | 0.50 | 0.19 | 1.28 |
| 40Hz | random masker | 0.15 | 0.06 | 0.37 |
| 50Hz | (Intercept) | 3097.80 | 1667.09 | 6760.91 |
| 50Hz | steady-state masker | 0.41 | 0.15 | 1.11 |
| 50Hz | random masker | 0.14 | 0.05 | 0.37 |
| 80Hz | (Intercept) | 3757.20 | 2122.84 | 7602.55 |
| 80Hz | steady-state masker | 0.41 | 0.16 | 1.01 |
| 80Hz | random masker | 0.07 | 0.03 | 0.17 |

**Table S14.**

**Experiment 2: ANOVA.** Analysis of variance for the predictor of playback condition for each negative binomial model, separately.

| Type II Analysis of Deviance | | | | |
|---|---|---|---|---|
| **model** | **predictor** | **LR Chi-square** | **df** | ***p*** |
| 4Hz | condition | 7.55 | 2 | 0.02 |
| 8Hz | condition | 0.31 | 2 | 0.86 |
| 16Hz | condition | 5.07 | 2 | 0.08 |
| 25Hz | condition | 7.55 | 2 | 0.02 |
| 33Hz | condition | 12.74 | 2 | <0.001 |
| 40Hz | condition | 15.16 | 2 | <0.001 |
| 50Hz | condition | 14.36 | 2 | <0.001 |
| 80Hz | condition | 27.63 | 2 | <0.001 |

**Table S15.**

**Number of temporally overlapping calls across both experiments in each condition**

| | Silent | Broadband masker (Steady-state masker) | High-freq masker | Random masker |
|---|---|---|---|---|
| Experiment 1 | 17,645 | 5,627 | 19,346 | - |
| Experiment 2 | 24,903 | 4,093 | - | 2,946 |

**Table S16.**

**Estimated marginal means for overlapping calls.** Estimated marginal mean differences in call incidence rates calculated on negative binomial models for overlapping calls for each modulation rate, separately.

| | | | Estimated Marginal Means | | | | | |
|---|---|---|---|---|---|---|---|---|
| **experiment** | **modulation** | **contrast** | **IRR** | **se** | **$CI_{lo}$** | **$CI_{hi}$** | **Z ratio** | ***p*** |
| 1 | 8Hz | silence / (broadband masker) | 2.40 | 1.28 | 0.66 | 8.65 | 1.63 | 0.31 |
| | 8Hz | silence / (high-freq masker) | 1.40 | 0.74 | 0.39 | 4.94 | 0.63 | 1.00 |
| | 8Hz | (broadband masker) / (high-freq masker) | 0.58 | 0.32 | 0.16 | 2.17 | -0.98 | 0.98 |
| | 15Hz | silence / (broadband masker) | 4.73 | 2.19 | 1.56 | 14.36 | 3.35 | <0.001 |
| | 15Hz | silence / (high-freq masker) | 0.37 | 0.17 | 0.12 | 1.11 | -2.17 | 0.09 |
| | 15Hz | (broadband masker) / (high-freq masker) | 0.08 | 0.04 | 0.03 | 0.24 | -5.44 | <0.001 |
| 2 | 4Hz | silence / (steady-state masker) | 4.58 | 3.04 | 0.94 | 22.45 | 2.29 | 0.07 |
| | 4Hz | silence / random masker | 2.33 | 1.33 | 0.60 | 9.11 | 1.49 | 0.41 |
| | 4Hz | (steady-state masker) / random masker | 0.51 | 0.34 | 0.10 | 2.58 | -1.00 | 0.96 |
| | 8Hz | silence / (steady-state masker) | 1.92 | 1.21 | 0.43 | 8.65 | 1.04 | 0.90 |
| | 8Hz | silence / random masker | 5.36 | 3.16 | 1.31 | 21.96 | 2.85 | 0.01 |
| | 8Hz | (steady-state masker) / random masker | 2.79 | 1.78 | 0.60 | 12.88 | 1.61 | 0.32 |
| | 16Hz | silence / (steady-state masker) | 2.21 | 1.16 | 0.63 | 7.77 | 1.52 | 0.39 |
| | 16Hz | silence / random masker | 6.56 | 3.32 | 1.95 | 22.02 | 3.72 | 0.00 |
| | 16Hz | (steady-state masker) / random masker | 2.96 | 1.60 | 0.81 | 10.83 | 2.00 | 0.14 |
| | 25Hz | silence / (steady-state masker) | 5.14 | 3.10 | 1.21 | 21.80 | 2.71 | 0.02 |
| | 25Hz | silence / random masker | 9.89 | 5.80 | 2.43 | 40.26 | 3.91 | 0.00 |
| | 25Hz | (steady-state masker) / random masker | 1.93 | 1.27 | 0.40 | 9.37 | 0.99 | 0.96 |
| | 33Hz | silence / (steady-state masker) | 5.72 | 3.07 | 1.58 | 20.69 | 3.24 | 0.00 |
| | 33Hz | silence / random masker | 14.70 | 7.47 | 4.35 | 49.66 | 5.28 | 0.00 |
| | 33Hz | (steady-state masker) / random masker | 2.57 | 1.50 | 0.64 | 10.41 | 1.62 | 0.32 |
| | 40Hz | silence / (steady-state masker) | 6.48 | 3.29 | 1.92 | 21.87 | 3.68 | 0.00 |
| | 40Hz | silence / random masker | 18.39 | 9.20 | 5.55 | 60.93 | 5.82 | 0.00 |
| | 40Hz | (steady-state masker) / random masker | 2.84 | 1.53 | 0.78 | 10.35 | 1.93 | 0.16 |
| | 50Hz | silence / (steady-state masker) | 11.27 | 6.43 | 2.88 | 44.16 | 4.25 | 0.00 |
| | 50Hz | silence / random masker | 27.70 | 15.51 | 7.26 | 105.80 | 5.93 | 0.00 |
| | 50Hz | (steady-state masker) / random masker | 2.46 | 1.45 | 0.60 | 10.12 | 1.52 | 0.38 |
| | 80Hz | silence / (steady-state masker) | 7.69 | 3.88 | 2.30 | 25.73 | 4.04 | 0.00 |
| | 80Hz | silence / random masker | 26.34 | 13.56 | 7.68 | 90.35 | 6.35 | 0.00 |
| | 80Hz | (steady-state masker) / random masker | 3.43 | 1.94 | 0.89 | 13.24 | 2.18 | 0.09 |

**Table S17.**

**Call timings relative to acoustic landmarks**

| Mean Call Onset Timings | | | | | | |
|---|---|---|---|---|---|---|
| | | | **time from peaks (ms)** | | **time to trough (ms)** | |
| **experiment** | **modulation** | **condition** | **median** | **iqr** | **median** | **iqr** |
| **1** | **All** | broadband masker | 37.7 | 27.9 | 10.1 | 1 |
| | 8Hz | broadband masker | 52.0 | 0.7 | 10.5 | 0.7 |
| | 15Hz | broadband masker | 24.1 | 0.7 | 9.6 | 0.7 |
| **2** | **All** | steady-state masker | 9.3 | 28.7 | 8.4 | 5.8 |
| | **All** | random masker | 9.6 | 26.5 | 9.0 | 8.4 |
| | 4Hz | steady-state masker | 83.7 | 2.8 | 41.3 | 2.8 |
| | 4Hz | random masker | 83.7 | 2.0 | 41.3 | 2.0 |
| | 8Hz | steady-state masker | 52.8 | 1.3 | 9.7 | 1.3 |
| | 8Hz | random masker | 46.1 | 1.4 | 16.4 | 1.4 |
| | 16Hz | steady-state masker | 25.8 | 0.9 | 5.0 | 0.9 |
| | 16Hz | random masker | 22.2 | 1.3 | 8.6 | 1.3 |
| | 25Hz | steady-state masker | 8.9 | 1.0 | 11.1 | 1.0 |
| | 25Hz | random masker | 12.2 | 1.7 | 7.8 | 1.7 |
| | 33Hz | steady-state masker | 10.5 | 2.9 | 4.3 | 2.9 |
| | 33Hz | random masker | 7.4 | 1.2 | 7.4 | 1.2 |
| | 40Hz | steady-state masker | 5.3 | 1.0 | 7.2 | 1.0 |
| | 40Hz | random masker | 4.4 | 2.5 | 8.1 | 2.5 |
| | 50Hz | steady-state masker | 0.5 | 1.4 | 9.5 | 1.4 |
| | 50Hz | random masker | -4.1 | 3.8 | 14.1 | 3.8 |
| | 80Hz | steady-state masker | 4.2 | 1.8 | 2.5 | 1.8 |
| | 80Hz | random masker | 0.5 | 5.8 | 6.2 | 5.8 |
